# S-acylation controls SARS-Cov-2 membrane lipid organization and enhances infectivity

**DOI:** 10.1101/2021.03.14.435299

**Authors:** Francisco S. Mesquita, Laurence Abrami, Oksana Sergeeva, Priscilla Turelli, Béatrice Kunz, Charlène Raclot, Jonathan Paz Montoya, Luciano A. Abriata, Matteo Dal Peraro, Didier Trono, Giovanni D’Angelo, F. Gisou van der Goot

**Author notes:** Corresponding Authors: F. Gisou van der Goot, Francisco S. Mesquita. These authors contributed equally (Francisco S. Mesquita, Laurence Abrami).

## Abstract

SARS-CoV-2 virions are surrounded by a lipid bilayer which contains membrane proteins such as Spike, responsible for target-cell binding and virus fusion, the envelope protein E and the accessory protein Orf3a. Here, we show that during SARS-CoV-2 infection, all three proteins become lipid modified, through action of the S-acyltransferase ZDHHC20. Particularly striking is the rapid acylation of Spike on 10 cytosolic cysteines within the ER and Golgi. Using a combination of computational, lipidomics and biochemical approaches, we show that this massive lipidation controls Spike biogenesis and degradation, and drives the formation of localized ordered cholesterol and sphingolipid rich lipid nanodomains, in the early Golgi where viral budding occurs. ZDHHC20-mediated acylation allows the formation of viruses with enhanced fusion capacity and overall infectivity. Our study points towards S-acylating enzymes and lipid biosynthesis enzymes as novel therapeutic anti-viral targets.

## INTRODUCTION

β-Coronaviruses (CoVs) belong to a family of large, positive-stranded RNA, enveloped viruses capable of infecting a wide range of hosts and with great zoonotic potential (V’kovski et al., 2020). Human-tropic species can cause from mild sub-clinical upper respiratory symptoms (i.e. common colds) to a life-threatening form of atypical pneumonia termed severe acute respiratory syndrome (SARS) (V’kovski et al., 2020; Wang et al., 2020), such as COVID-19 caused by SARS-CoV-2 (SARS coronavirus type 2) (Bojkova et al., 2020; Gordon et al., 2020). Despite the impressive advances in understanding SARS-CoV-2 over the last year, a precise molecular understanding of the biogenesis of the virus and of the factors that govern infectivity is still lacking. CoVs are surrounded by a membrane in which at least 3 structural proteins are embedded: Spike (S), Membrane (M), and Envelope (E) (Boson et al., 2021; Siu et al., 2008; V’kovski et al., 2020) as well as accessory factors such as Orf3a (Ito et al., 2005; Tan et al., 2004). Viral assembly requires E and M (Boson et al., 2021; Siu et al., 2008) but not Spike (Siu et al., 2008). Spike is however essential for infectivity as it mediates attachment to the host cell receptors, angiotensin-converting enzyme 2 (ACE2) for SARS-CoV-2 (Hoffmann et al., 2020b; Letko et al., 2020), and triggers the fusion between viral and target cell membranes, to release the viral genome into the cytoplasm. It is therefore the target of most vaccine strategies (Dai and Gao, 2020; Hoffmann et al., 2020a).

Spike, E and M are synthesized by ribosomes associated to the ER, where folding and quaternary assembly occurs, such as trimerization of Spike. They may also undergo post-translation modifications such as N-glycosylation on their luminal domains (Watanabe et al., 2020) and S-acylation (commonly referred to as S-palmitoylation) on their cytosolic domains. Both S and E have been reported to undergo S-acylation in the case of SARS-CoV-1, mouse hepatitis virus (MHV) and other CoVs (Boscarino et al., 2008; Lopez et al., 2008; McBride and Machamer, 2010; Petit et al., 2007; Thorp et al., 2006; Tseng et al., 2014). S-acylation, which consists in the covalent attachment of medium chain fatty acids (frequently palmitate, C16) to the sulphur atom of cytosolic cysteines, is very frequent in mammalian cells and considered a hallmark of viral envelope proteins (Gadalla and Veit, 2020; Schmidt, 1982). S-acylation of Spike has been proposed to mediate its association to lipid microdomains, promote syncytia formation upon transfection into cells and reduce infectivity of MHV and of pseudotyped particles (McBride and Machamer, 2010; Nguyen et al., 2020; Sanders et al., 2020; Thorp et al., 2006).

Here, we present a comprehensive study of the role of S-acylation in the SARS-CoV-2 infection cycle, including the detailed characterisation of Spike acylation, the identification of additional acylated viral proteins and host enzymes involved, and the importance of acylation in viral biogenesis and infection. We identify ZDHHC20 as the main acyltransferase responsible for the lipidation of SARS-CoV-2 Spike, E and Orf3a. We demonstrated that Spike acylation plays several roles, promoting biogenesis by protecting from premature targeting to ER degradation pathway, increasing the Spike half-life, and controlling the lipid composition of its immediate membrane environment. Lipidomic analyses reveal the unique lipid composition of SARS-CoV-2 virions, enriched in specific early Golgi sphingolipids, quite distinct from that of viruses budding from the plasma membrane. Finally, we show that S-acylation and sphingolipid metabolism promote SARS-CoV-2 infection, pointing towards S-acylation and lipid biogenesis related enzymes as novel potential anti-viral targets.

## RESULT and DISCUSSION

### Acylation of SARS CoV-2 viral proteins by ZDHHC20

Consistent with previous findings on SARS-CoV-1 (McBride and Machamer, 2010), SARS-CoV-2 Spike protein can undergo S-acylation when ectopically expressed in Hela cells. We showed that this also occurs during infection of monkey kidney Vero E6 cells with SARS-Cov-2 (Figure 1A-D), when a large proportion of infected cells displayed high levels of Spike protein (Figures S1AB). We used two established assays to monitor protein acylation. The <u>Acyl</u>-<u>R</u>esin <u>A</u>ssisted <u>C</u>apture assay (Acylrac) allows the capture of S-acylated proteins using thiol-reactive Sepharose beads. These can capture cysteine thiol groups, which are exposed by a cleavage step with hydroxylamine, that breaks the thioester bond between target proteins and the fatty acid moiety (Figure 1AC). The second assay consists in incubating cells with radioactive ^3^H-palmitate, and monitoring the incorporation by Spike using immunoprecipitation and autoradiographic analysis (Figure 1BD). Several Spike bands can be observed because to become fusion competent, Spike undergoes two sequential cleavage steps, mediated by host proteases, at the S1-S2 site and subsequently the S2’ site. The S2 fragment contains the membrane fusion subunit and the transmembrane anchor (Hoffmann et al., 2020a, 2020b). Western blot analysis of total cell extracts also showed a smaller C-terminal fragment (Figure 1AC). Acylation of all Spike fragments could be readily detected (Figure 1A-D) for both SARS-CoV-2 WT Spike originating from the initial Wuhan isolate, and the D614G variant.

**FIGURE 1:**
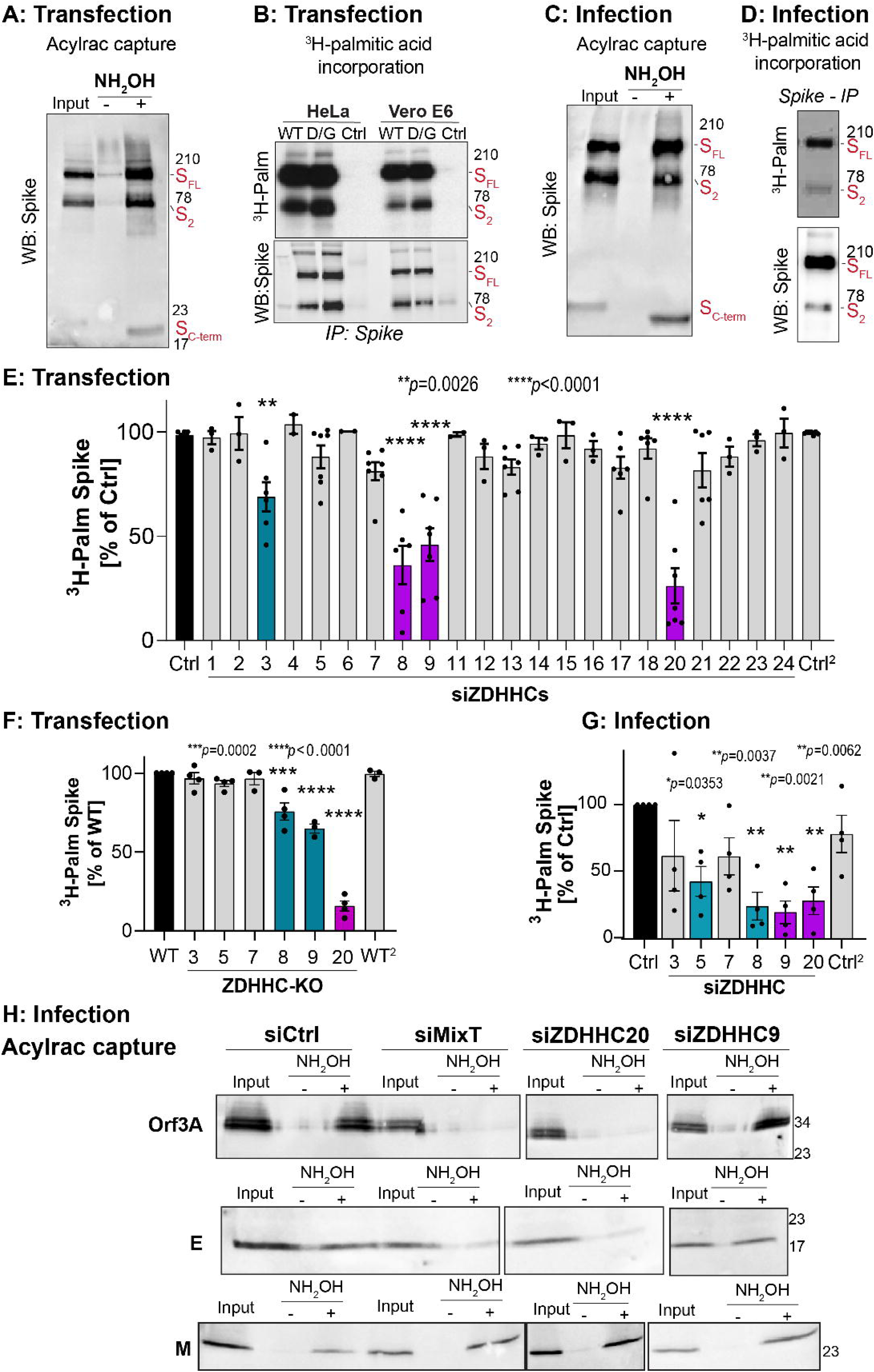
Identification of the Spike modifying ZDHHC acyltransferases. **See alos Figure S1. A.** Vero E6 cells expressing Spike WT (24 h) were lysed and S-acylated proteins detected using Acylrac capture assay. Cell lysate fractions (input), S-palmitoylated proteins (detected after hydroxylamine treatment, +NH_2_OH) and control fractions (-NH_2_OH) were analyzed by western blot against Spike. Three bands were detected: Spike Full length (FL), the S2 and the C-terminal fragment (C-term). **B.** Hela or Vero E6 cells expressing WT or D614G (D/G) mutant Spike were metabolically labeled for 2 h at 37°C with ^3^H-palmitic acid. Spike was immunoprecipitated and samples analyzed by western blot against Spike or parallel autoradiography (^3^H-Palm). **CD.** Vero E6 cells infected with SARS-CoV-2, (MOI≈0.1, 24 h) were: **C.** processed for Acylrac or **D** metabolically labeled as in B, but throughout the 24 h of infection, when Spike S-palmitoylation was analyzed. **E.** Analysis of Spike-incorporated radioactivity in Hela cells co-transfected with individual siRNAs targeting all human ZDHHCs or control siRNA (Ctrl) for 72 h and plasmids expressing Spike (48 to 72 h). Incorporated radioactivity, as described in B for Spike Full length (FL) was quantified using the Typhoon Imager and normalized to control cells. Results are mean ± SEM, and each dot represents an individual independent experiment (n ≥ 3). **F.** Analysis of Spike-incorporated radioactivity (as in E) in HAP-1 cells knockout for the indicated ZDHHCs, and ectopically expressing Spike (24 to 48 h). **G.** Analysis of Spike-incorporated radioactivity in Vero E6 cells transfected with siRNA targeting individual ZDHHCs or control siRNA (siCtrl) and infected and processed as in C, D. Results are mean±SEM, n = 4 independent experiments. For **E**, **F** and **G** two independent Control samples were analyse (Ctr^2^ and WT^2^) and *p*-values were obtained by One-way ANOVA with Dunnet’s multiple comparison test. **H** Acylrac analysis in Vero E6 cells transfected with individual siZDHHC9, siZDHHC20, pool siMixT (ZDHHC8, ZDHHC9 or ZDHHC20) or control siRNAs (siCtrl), and infected with SARS-CoV-2, (MOI≈0.1, 24 h). Samples were analyzed by Western blot with antibodies against SARS-CoV-1/2 proteins Orf3a, E and M. An effect of silencing ZDHHC5 on ^3^H-palmitate incorporation (Figure 1F) could not be interpreted because it also triggered a drastic drop in Spike expression **(Figure S1F)**, for reasons that remain to be determined.

Although S-acylation of enveloped virus proteins has been known since decades, the mechanisms and mediators remain ill-defined. S-acylation is catalysed by a family of 23 protein acyltransferases defined by the presence of a conserved zinc-finger domain with a DHHC motif in their catalytic site (ZDHHCs) (Zaballa and Goot, 2018). To identify the acyltransferases responsible for the modification of Spike, we individually depleted the 23 ZDHHC enzymes by RNA interference in Hela cells (ZDHHC1 to 24, ZDHHC10 does not exist). Downregulation of three family members, ZDHHC8, 9 and 20, led to a >50% drop in ^3^H-palmitate incorporation into ectopically expressed Spike (Figure 1E, S1C). A subset screen, carried in near-haploid HAP-1 cells knocked out for individual ZDHHCs, confirmed these findings and indicated a principal role for ZDHHC20 (Figure 1F, S1D). Spike acylation upon depletion of the individual ZDHHC8, 9 or 20 could be restore by expression of the correspondent siRNA-resistant myc-tagged ZDHHC enzyme, but not a random acyltransferase (Figure S1E). In line with our findings in HAP1 cells, ectopic expression of ZDHHC20 could also fully rescue Spike acylation upon depletion of ZDHHC8 or ZDHHC9, further supporting a prominent role for ZDHHC20 (Figure S1E).

We next tested the involvement of the identified ZDHHC enzymes during infection in Vero E6 cells, using monkey specific siRNAs. Expression of ZDHHCs varies significantly between cell types (Figure S1G), with ZDHHC20 being more abundant in Vero E6 cells when compared to the human cell lines used in this study (Figure S1D). We first targeted ZDHHC8, 9, and 20 using an siRNA mix (MixT). Acylated Spike could no longer be retrieved using the Acylrac capture assay (Figure S1E). We then silenced the enzymes individually, and included additional ZDHHCs for comparison (Figure 7G and S7G). Unfortunately, efficient depletion (below 50%) of monkey ZDHHC8 also led to a decrease in monkey ZDHHC20 mRNA in Vero cells, an effect we did not observed in human cell lines (Figure S1H). We therefore omitted ZDHHC8 from further analysis in Vero cells, but maintained the use of MixT. Knocking down ZDHHC9 and 20 led to a strong decrease in S-acylation of Spike during infection (Figure 1G).

Since Spike is not the only acylated SARS-CoV-2 protein, we tested M, E and Orf3a, which all have cytosolic cysteines (Figure S1I) (Boscarino et al., 2008; Issa et al., 2020, p. 3). Acylrac analysis indicated that all three proteins are acylated during infection. Interestingly, E and Orf3A, but not M, could no longer be retrieved when cells were pre-treated with siMixT or siZDHHC20 (Figure 1H). Altogether, these observations show that Spike is a substrate for ZDHHC9 and ZDHHC20 (monkey and human), and ZDHHC8 (human), and that ZDHHC20 also modifies E and Orf3a.

### Rapid and extensive acylation during Spike biogenesis

*Coronaviridae* Spike proteins contain a conserved cysteine-rich stretch of amino acids in their cytosolic tail (Figure 2A) (Gelhaus et al., 2014; Thorp et al., 2006). SARS-CoV-2 Spike has 10 cysteines within its first 20 cytosolic amino acids (Figure 2A) and was recently highlighted as one of the most cysteine-rich proteins encoded by animal viruses (Sanders et al., 2020). To study the relative importance of these residues, we generated four mutants changing groups of 2-3 adjacent cysteines to alanine (Figure 2A). In transfected cells, mutation of the two most membrane-proximal cysteines (group I) led to an 80% drop in ^3^H-palmitate incorporation into the Spike protein (Figure 2BC), as previously observed for SARS-CoV-1 Spike (Petit et al., 2007). Mutation of the next three cysteines (group II) also had a major, albeit, lower effect with a ≈40% drop in ^3^H-palmitate incorporation (Figure 2BC). Mutating cysteines from both groups I and II had an impact almost equivalent to that of changing all 10 to alanine (Figure 2BC). Yet, cysteines from groups III and IV also undergo S-acylation, since mutating both groups alone or in conjunction with group II still significantly reduced ^3^H-palmitate incorporation (Figure 2BC). Altogether this analysis indicates that all 10 cysteines are targets of S-acylation and lipidation of those closest to the membrane is required for the others to be modified, possibly because it brings the more distal cysteines closer to the membrane, and therefore more accessible to the ZDHHC active site.

**FIGURE 2:**
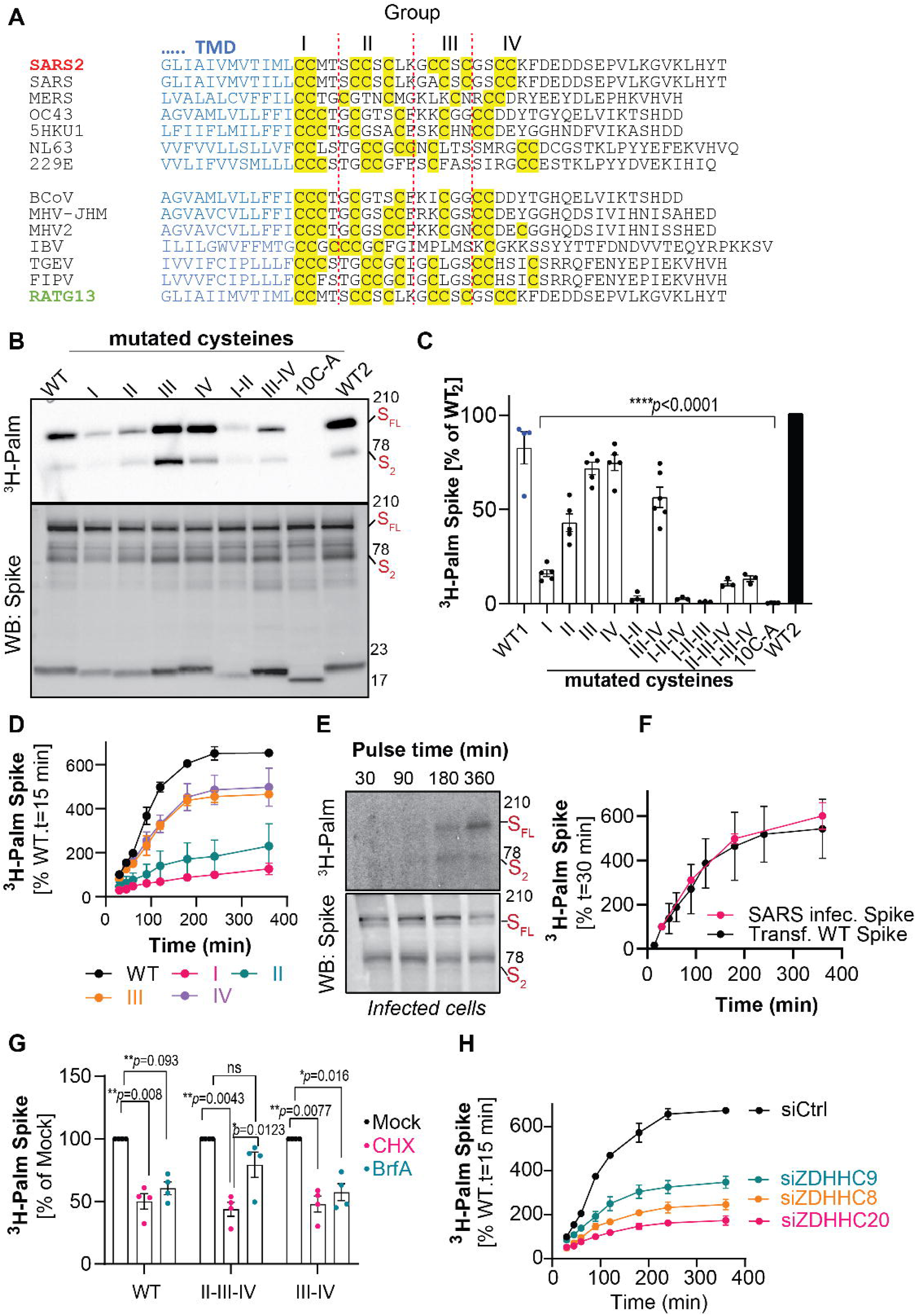
ZDHHC20 initiates acylation on the two juxtamembranous cysteines. **See alos Figure S2. A.** Alignment of the juxtamembranenous regions of various Spike proteins (top-human specific, bottom various animal species). Transmembrane domains are shown in blue and yellow highlights depict cysteine residues that were split into 4 groups: I, II, III, IV. The protein accession numbers are: SARS2 (A0A7D6HN98), SARS (P59594), MERS (K9N5Q8), OC43 (P36334), 5HKU1 (A0A451ERM8), NL63 (H9EJ93), 229E (A0A1YOEV30), BCov (P25191), MHV-JHM (P11225), MHV2 (Q77NQ7), IBV (P11223), TGEV (P07946), FIPV (P10033), RATG13 (A0A6B9WHD3). **B.C.** Hela cells were transfected 24 h with the indicated HA-tagged Spike variants (WT, or mutants in which, individual or combined cysteine groups (depicted in A) were replaced by alanine; all cysteines were mutated to for 10C-A mutant). Cells were metabolically labeled (2 h) with ^3^H-palmitic acid and HA-Spike-immunoprecipitation (IP) fractions analyzed by western blot with HA-HRP antibodies or autoradiography (^3^H-Palm). **C.** Levels of incorporated radioactivity for Spike Full length (FL) and S2 were quantified using the Typhoon Imager and normalized to one of two independent WT samples (WT2). Results are mean ± SEM, and each dot represents an independent experiment (n ≥ 4). **D.** Levels of Spike incorporated radioactivity analyzed in IP-Spike fractions from Hela cells expressing the indicated HA-tagged Spike variants metabolically labeled for different times with ^3^H-palmitic acid. Values for WT at 15 min were set to 100% and used as reference. Results are mean ± SD, n = 3 independent experiments. **E.** Vero E6 cells were infected 4 h with SARS-CoV-2, (MOI≈0.5-1). Cells were metabolically labeled with ^3^H-palmitic acid for different times, processed as in B. **F.** Spike Incorporated radioactivity quantified as in **C.** was normalized to the increasing levels of IP-Spike. Values at 30 min were set to 100% and used as reference. Results are mean ± SD, n = 3 independent experiments. The ^3^H-palmitic acid incorporation by untagged Spike in HeLa cells (Transf –WT Spike) was quantified in parallel and displayed for comparison. **G.** Quantification of Spike incorporated radioactivity in Hela cells expressing the indicated Spike variants pre-treated (1 h) with serum-free medium with Cycloheximide (CHX), Brefeldin A (BrfA) or Mock-medium before ^3^H-palmitic acid metabolic labelling. Values for Mock treated cells were set to 100% for each Spike mutant. Results are mean ± SEM, and each dot represents an independent experiment (n = 4). **H.** Hela cells transfected (72 h) with siRNAs targeting the indicated ZDHHCs or control siRNA (Ctrl) and plasmids expressing Spike WT (24 to 48 h) were metabolically labeled and analyzed as in **D**. Results are mean ± SD, n = 3 independent experiments. *p* values were obtained by **C** One-way or **G** Two-way ANOVA with Dunnet’s or Tukey’s multiple comparison tests respectively.

To gain more insight into the dynamics of this process, we measured ^3^H-palmitate incorporation kinetics in transfected HeLa cells. Incorporation of ^3^H-palmitate into WT Spike was rapid, reaching a plateau within about 3 h, in great contrast with group I- and group II-mutated variants, where it was very slow and limited (Figure 2D and S2A). Mutation of groups III and IV cysteines had no major effect on S-palmitoylation kinetics but resulted in lower final levels of incorporation, consistent with the removal of 3, or 2 acceptor amino acids, respectively (Figure 2D). Thus, membrane proximal cysteines play a more influential role in Spike S-acylation than their more distal counterparts.

We also evaluated ^3^H-palmitate incorporation into Spike during a single-round of infection in Vero E6 cells. We used a higher MOI for infection (≈0.5 to 1) and added radioactive ^3^H-palmitate 4 h post-viral inoculation, when Spike started being detected by flow cytometry and significant viral RNA replication was observed by QPCR (Figure S2A-C). Remarkably, the kinetics of Spike ^3^H-palmitate incorporation were essentially identical for infected and transfected cells (Figure 2EF and S2D).

We next investigated where during viral biogenesis Spike acylation occurs. After synthesis, folding and assembly, Spike trimers are transported to the ER-Golgi intermediate compartment (ERGIC) where viral assembly and budding take place (de Haan and Rottier, 2005; Klein et al., 2020; Stertz et al., 2007). Assembled virions are subsequently transported to the plasma membrane to be released through exocytic mechanisms involving the Golgi apparatus and/or lysosomal compartments (Ghosh et al., 2020). To determine if Spike acylation occurs during protein biogenesis, and/or after transport from the ER to the Golgi, we monitored ^3^H-palmitate incorporation under the effects of the protein synthesis inhibitor cycloheximide (CHX), and the Golgi-disrupting drug Brefeldin A. Incorporation into WT Spike was decreased by ≈50% when cells were treated with either CHX or Brefeldin A (Figure 2G), suggesting that S-palmitoylation is initiated shortly after protein synthesis and continues once the protein has exited the ER. This is consistent with previous reports on SARS-CoV-1 and MHV Spike proteins indicating that their palmitoylated versions were present in both pre- and post-Golgi compartments (McBride and Machamer, 2010; van Berlo et al., 1987). We next determined the effect of the drugs on the S-palmitoylation of two additional mutants, bearing respectively only the first 2 (groups II, III and IV mutated) or 5 (groups III and IV mutated) most membrane-proximal cysteines (Figure 2G). With the former mutant Spike, sensitivity to brefeldin A was lost, suggesting that acylation of the 2 juxta-membrane cysteines occurs in the ER, whereas the latter had a profile similar to that of WT indicating that more distal cysteines are modified in downstream compartments.

We also monitored the kinetics of ^3^H-palmitate incorporation when silencing the Spike-modifying ZDHHC enzymes. ZDHHC20 has a broad distribution throughout the entire biosynthetic pathway from the ER to the plasma membrane (Figure S2E). ZDHHC9 expression appears to be more restricted to the ER, while ZDHHC8 is more concentrated in the perinuclear-Golgi region (Figure S2E). Silencing of any of the three enzymes had a drastic effect on ^3^H-palmitate incorporation, but silencing ZDHHC20 caused the earliest and most dramatic effect (Figure 2H), phenocopying the group I cysteine mutant (Figure 2D).

Altogether, these results indicate that following synthesis, Spike can undergo acylation on its two most juxtamembranous cysteines in the ER by ZDHHC20, a step required for the modification of the more distal cysteines by ZDHHC20, 8 or 9 in the ER and Golgi.

The above experiments, whether by Acylrac or ^3^H-palmitate incorporation, demonstrate that acylation occurs but do not indicate whether only a small percentage or all Spike molecules undergo the modification. A variant of Acylrac has therefore been developed where after removal of the acyl chains using hydroxylamine, the newly freed up cysteines are labelled with PEG (5 kDa), leading to molecular weight shifts (Howie et al., 2014; Percher et al., 2017). For small proteins, each PEGylation reaction leads to an approximate 5 kDa shift in the SDS-PAGE migration pattern. When proteins are large, and the number of acylation sites is important, molecular weight shifts are less predictable. When Spike was transfected into Vero E 6 cells, we observed a weak high molecular weight smear, but the full length or S2 Spike bands remained essentially unaltered (Figure 3A, left panel). In contrast, PEGylation of proteins from infected cells led to the disappearance of the S2 band (Figure 3A, right panel and 3B). It is unclear where exactly the S2 band shifted to. It could be the band migrating slightly higher than the full-length form. Extensive PEGylation may also hinder the full-length Spike from entering into the SDS gels. The disappearance of the S2 band nevertheless indicates that all cleaved Spike proteins were highly acylated during infection. A kinetic analysis of PEGylation indicated that Spike S-acylation occurred as early as 2 h after viral inoculation (Figure S3A), consistent with the rapid dynamics observed in our previous ^3^H-palmitate incorporation experiments (Figure 2). We also performed PEGylation on the virion containing supernatants of infected Vero E6 cells and on isolated virions, observing similar shifts (Figure 3CD). The PEG induced band shifts in S2 were abolished when ZDHHC enzymes were silenced with MixT or ZDHHC20 siRNA (Figure 3B-D and S3B). Correspondingly capturing of acylated Spike from supernatants of infected cells was also abrogated when cells were silenced with MixT (Figure S3C). The PEGylation analysis, combined with the ^3^H-palmitate incorporation experiments, indicate that the entire Spike population produced by infected cells undergoes massive and rapid acylation, apparently on all 10 cysteines.

**FIGURE 3:**
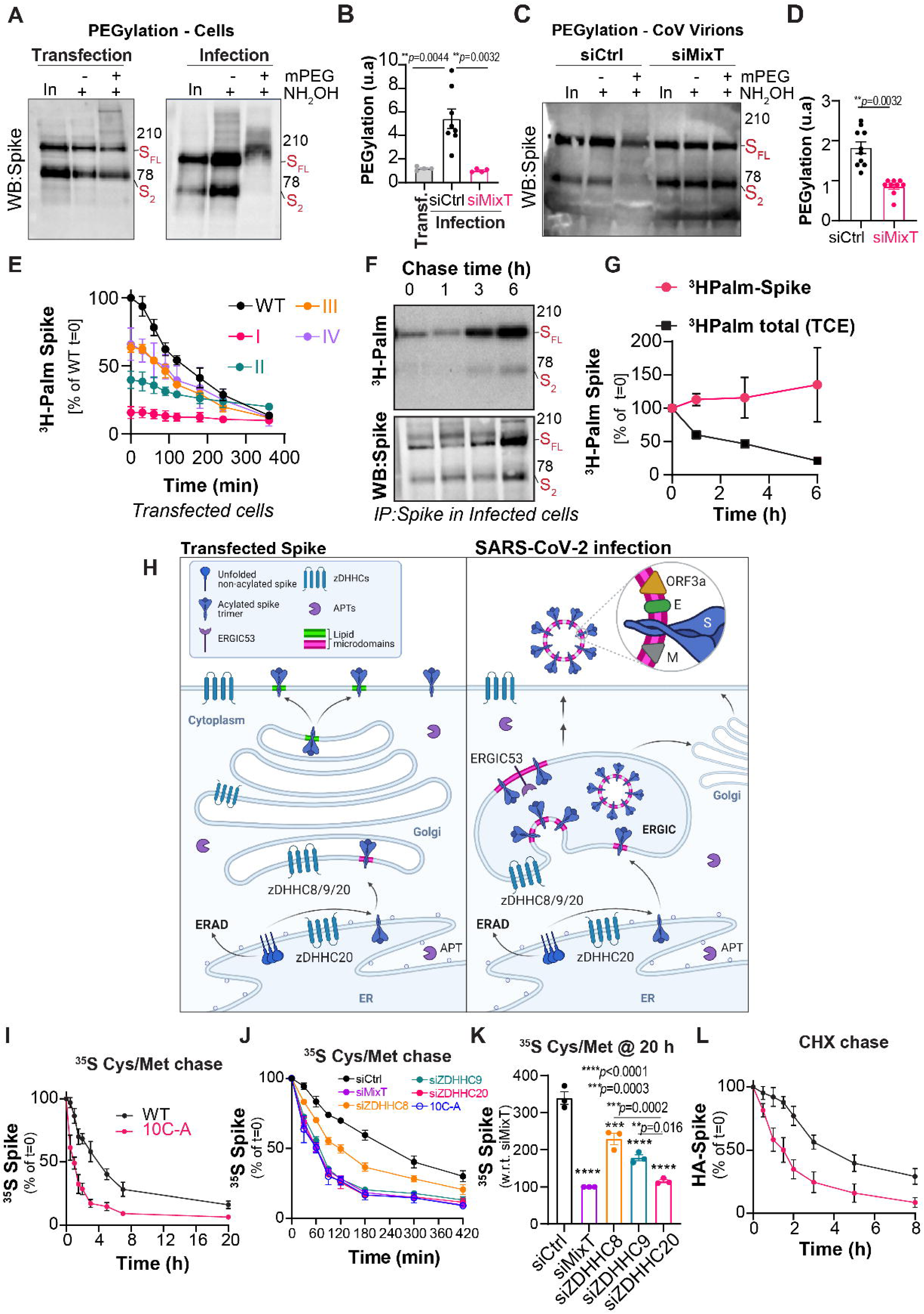
Acylation of Spike during infection is extensive and stable. **See alos Figure S3. A.** Acyl-PEG-exchange analysis (see experimental procedures) in lysates from Vero E6 cells expressing WT untagged Spike (24 h, left) or infected with SARS-CoV-2, MOI=0.1, 24 h (right). S-acylated proteins were labelled with PEG-5KDa (+mPEG) following hydroxylamine treatment (NH_2_OH) and analyzed with control non-labelled (-mPEG), and input cell lysate fractions (input) by western blot against Spike. **B.** Quantification of Spike-PEG migration shift (see experimental procedures) for the samples described in A (Transfected Spike – Transf) and for Spike from Vero E6 cells transfected (72 h) with Control or pool siRNA oligos targeting siZDHHC8, ZDHHC9 and ZDHHC20 (siMixT) and infected with SARS-CoV-2, MOI=0.1 24 h. Results are mean ± SEM, and each dot represents an independent experiment (n ≥ 4). **C and G.** Acyl-PEG-exchange analysis and quantification of pre-cleared, filtered supernatants from samples infected as in B. Results are mean ± SEM, and each dot represents an independent experiment of n=10 independent experiments. **E.** Hela cells expressing (24 h) the indicated HA-tagged Spike variants (depicted in Figure 2A) were metabolically labeled for a 2 h pulse with ^3^H-palmitic acid, and chased in complete medium for the indicated time-points. Incorporated radioactivity in IP-HA-Spike fraction was quantified and values for WT at T0 min set to 100% as reference. Results are mean ± SD, n = 3 independent experiments. **F** Vero E6 cells infected with SARS-CoV-2, (MOI≈0.5-1) were metabolically labeled with ^3^H-palmitic acid from 1 to 14 h post viral inoculation. Cells were chased in fresh complete medium and incorporated radioactivity in IP-Spike fractions analyzed by autoradiography and parallel western blot analysis. Values for Spike-incorporated radioactivity were normalized to the levels of IP-Spike protein and set to 100% for T=0 min samples. A fraction of total cell extracts (TCE) was used to quantify total incorporated radioactivity (See Figure S3F). Results are mean ± SD, n = 3 independent experiments **H.** Model of S-acylation-dependent control of Spike cellular distribution and membrane lipid organization during Spike ectopic expression or SARS-CoV-2 infection. Created with BioRender.com. **I.** Hela cells expressing HA-tagged Spike (WT or 10C-A mutant) were metabolically labelled with ^35^S-Met/Cys for 4 h, and chased in complete medium for the indicated time-points. The levels of ^35^S-Met/Cys radioactivity incorporated into IP-HA-Spike fractions were quantified and set to 100% for T=0 min for each sample. Results are mean ± SD, n = 4 independent experiments. **I.** Quantification of ^35^S-Met/Cys radioactivity incorporated into IP-HA-Spike fractions from HeLa cells expressing HA-tagged Spike (WT or 10C-A) and transfected (72 h) with Control siRNA oligos targeting siZDHHC8, ZDHHC9, ZDHHC20 or all combined-MixT. Values were set to 100% for T0 min for each sample. Results are mean ±SD, n = 3 independent experiments **J.** The 35S-Met/Cys radioactivity incorporated into IP-HA-Spike fractions following 20 h of chase were normalized to 100% for siMixT and displayed as mean ±SD, for n = 3 independent experiments **K.** HeLa cells expressing HA-tagged Spike (WT or 10C-A) were incubated in serum free medium with cycloheximide (CHX) throughout time. At indicated time points total cell extracts were analyzed by western blot against Spike and the levels of Spike Full length (FL) expressed relative to T=0 set as 100%. Results are mean ±SD, n = 4 independent experiments. All p values were obtained by B One-way ANOVA with Tukey’s or (J) Dunnet’s multiple comparison test or G unpaired student’s T-test.

### Spike acylation is stable during infection

S-acylation can be reversed by the action of deacylating enzymes, the acyl protein thioesterases (APTs) (Zaballa and Goot, 2018). Using a metabolic ^3^H-palmitate pulse-and-chase approach in transfected HeLa cells, we found that WT Spike lost its palmitate with a half-live of about 3 h, as did group III and IV mutants (Figure 3E and S3D), and that this process could be blocked using the general APT inhibitor Palmostatin B (Figure S3E). These data prompted us to test if S-acylation of Spike is also dynamic in infected cells.

We performed a similar ^3^H-palmitate decay analysis during SARS-CoV-2 infection of Vero E6 cells. ^3^H-Palmitate was added 14 h post-inoculation for 2 h, and cells were further incubated in label-free medium. While depalmitoylation of cellular proteins (detected by autoradiography of total cell extracts) was readily observed (Figure S3F and 3G), there was no significant loss of palmitate from Spike (Figure 3FG). This observation suggests that Spike deacylation barely occurs during infection of Vero E6 cells. The difference with depalmitoylation of Spike during transfection could be due to rapid segregation of Spike from the acyl thioesterases during formation of virions in the early Golgi, while the Spike cysteines remain cytosolically exposed to acyl thioesterases in transfected cells (model in Fig. 3H).

### S-acylation promotes the biogenesis of Spike and reduces its turnover rate

S-acylation has been reported to modify the turnover rate of proteins (Zaballa and Goot, 2018). We therefore tested whether this was the case for Spike. We used two different methods, ^35^S Cys/Met metabolic pulse-chase experiments, which monitor the evolution of a protein starting from its synthesis, and cycloheximide chase, which monitors the degradation of fully folded mature proteins. The ^35^S Cys/Met pulse period was chosen based on the plateau of palmitate incorporation, i.e. 4 h (Figure 2D). The decay in the ^35^S signal observed for WT Spike indicated an apparent half-life of ≈4 h, while that observed for the 10C-A cysteine-less mutant was ≈1 h (Figure 3I and S3G). The accelerated early decay for the 10C-A mutant suggests that this derivative is rapidly targeted to degradation in the ER, whereas acylation protects the WT protein from this premature process. The targeting to degradation is indicative of inefficient folding, recognized by the ER quality control machinery. A previous study did not reveal an effect of SARS-Cov-1 Spike S-acylation on protein degradation (McBride and Machamer, 2010). In these experiments, the ^35^S Cys/Met metabolic labelling time was short, 20 min, but we know now this is insufficient to significantly populate the acylated Spike species (Figure 2D) (Abrami et al., 2019). Acceleration of Spike degradation following its synthesis was also observed when silencing the ZDHHC enzymes (Figure 3J). Silencing ZDHHC9 or 20 was comparable to siMixT or mutating all cysteines, while silencing of ZDHHC8 had a less pronounced effect (Figure 3K). Comparison of the total levels ^35^S Cys/Met-labelled Spike at 20 h of chase showed that acylation, primary by ZDHHC20, leads to a 3-fold increase in total Spike (Figure 3K). Thus, acylation is crucial for efficient Spike biogenesis. In addition, through the cyloheximide chase approach, we found that S-acylation also slows down the turnover of mature, fully folded, Spike (Figure 3L and S3H).

Therefore, by increasing the flux of newly synthesized Spike through the ER quality control gateway and by slowing down the turnover of the fully folded protein, S-acylation dually increases the levels of Spike available for virus assembly.

### Spike S-acylation modifies the organization of the lipid bilayer

Following synthesis, Spike assembles into trimers (Delmas and Laude, 1990), which for SARS-CoV-2 can each bear 30 fatty acids. Cryo-electron microscopy studies indicate the presence of 25 to 130 Spike trimers per virion (Bar-On et al., 2020; Ke et al., 2020; Klein et al., 2020). With a 80-90 nm virus diameter and an average headgroup surface of 70 Å^2^ per phospholipid, a “back-of-the-envelop” calculation indicates that the acyl chains attached to Spike could represent 2-12% of the lipid molecules of the inner leaflet of the viral membrane, if randomly distributed.

Given this very significant contribution, we analyzed the possible impact of Spike acylation on the surrounding lipid organization, using coarse-grained (CG) molecular dynamics (MD) simulations. We compared the behavior of palmitoylated and non-palmitoylated Spike in a model membrane. We set up a membrane composed of 50% dipalmitoylphosphatidylcholine (DPPC), 30% dilinoleylphosphatidylcholine (DLiPC) and 20% cholesterol (CHOL), and parameterized it using the MARTINI CG force field, which is known to capture lipid nanodomain formation (Lorent et al., 2017; Risselada and Marrink, 2008). We first reproduced the results of Lorent et al., 2017 of microdomain formation (a liquid-disordered DLiPC-rich phase and a liquid-ordered DPPC+CHOL-rich phase) within ≈1 µs when starting from randomly distributed lipids. We subsequently prepared two systems, again with lipids distributed randomly, and introduced the transmembrane domain (TMD) of Spike followed by the cytosolic tail residues (residues 1201 to the end, Uniprot P0DTC2), either fully S-palmitoylated or not at all. We ran each system 10 times for ≈3.7-3.8 µs in CG MD simulations, starting from different random seeds (Figure 4A). Analysis of the MD simulations led to two predictions. While the cytoplasmic tails of the protomers were initially introduced in an extended form, acylation of all cysteine residues led to the collapse of the cytosolic tails, bringing the palmitate moieties in close proximity of the TMD and, leading to a ≈9-fold local increase in the acyl chain concentration (Figure 4AB). Such compaction is not an artefact of the coarse-grained description of the protein, as it was also observed in atomistic MD simulations using the CHARMM36m force field (Figure 4B). Spike preferentially associated with the disordered phase when unmodified, but rather partitioned into the ordered phase when S-palmitoylated. This preference was dictated by the cytosolic half of the TMD (Figure 4AB and S4A). In contrast, the luminal half of the TMD interacted preferentially with the disordered phase, irrespective of Spike acylation (Figure 4B and S4A). These simulations raised the interesting possibility, not tested here, that acylated Spike could drive trans-bilayer lipid asymmetry.

**FIGURE 4:**
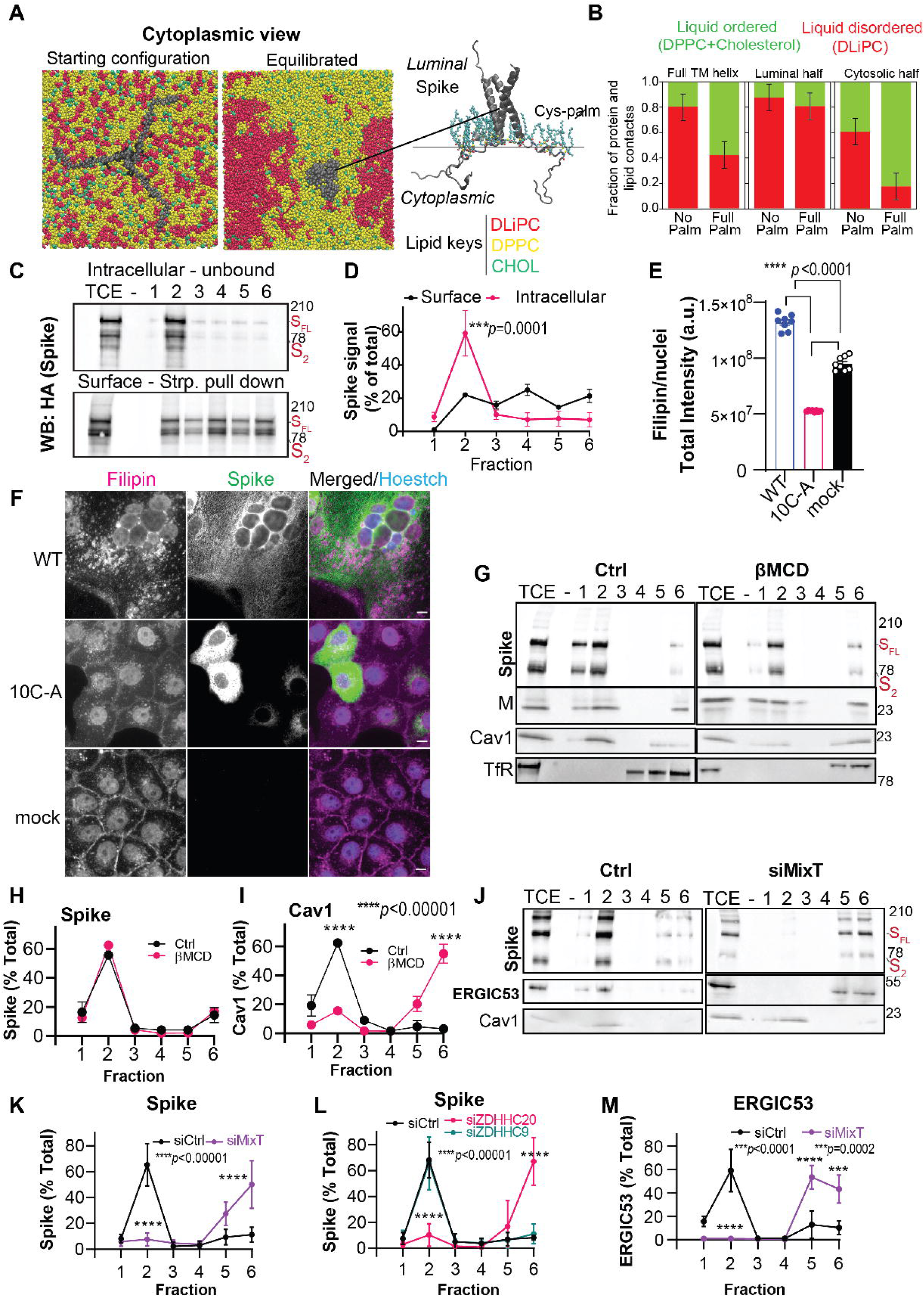
S-Acylation controls the lipid environment of Spike. **See alos Figure S4. A.** Coarse-grained (CG) model of Spike’s transmembrane (TM) helix + C-terminal region (residues 1201-end in Uniprot P0DTC2) assembled in trimeric form and inserted into a DPPC-(yellow)-DLiPC-(red)-cholesterol-(green) membrane model used in MD simulations. **Left**, starting MD conditions with random lipid positions and stretched cytoplasmic tail of Spike; and **center**, same view after 3.75 μs of CG MD simulation of the S-palmitoylated S protein. A representative end-state extracted from 10 CG MD replicas shows phase separation and Spike partitioning. In **right** the same model after 2 μs of atomistic MD simulation in the palmitoylated form in a POPC membrane **B.** Quantification of contacts established between the full TM helix, the luminal half, or the cytosolic half, and each lipid phase, after all phases were equilibrated. Results are averages from 10 simulations, with standard deviations. **C.** Vero E6 cells expressing HA-tagged Spike WT (WT) were surface labelled with biotin and processed for fractionation and DRM isolation (see experimental procedures). Surface biotinylated proteins (Surface-Strp. pull down) were precipitated from each fraction with streptavidin beads and compared to unbound fractions (intracellular-unbound) by western blot against Spike-HA. **D**. Quantification of Spike-HA (all forms) in each fraction relative to the total signal in all 6 fractions. Results are mean ± SD, n = 3 independent experiments. **EF.** Vero E6 cells expressing HA-tagged Spike (WT, or 10C-A mutant) for 24 h were fixed, immunolabeled for HA and stained with Filipin. Images were acquired and quantified by high-throughput automated microscopy and **(E)** the total levels of filipin staining within HA-positive-cells (or per cell for mock sample) quantified and divided by the number of nuclei. The data was averaged for 49 frames over 8 different wells per condition. Each dot represents one independent well, and results are mean ± SEM. Equivalent results were obtained for three independent assays **G.** Vero E6 cells infected with SARS-CoV-2 24 h, MOI≈0.1 were incubated with beta-methylcyclodextrin (βMCD) or not (Ctrl) for 30 min. Cells were processed as in **C** and analyzed by western against Spike, The DRM marker Caveolin1 (Cav1) was used as positive control and Transferrin Receptor (TfR) as negative control. **H**.**I** Quantification of Spike (all forms) (**H**) and Caveolin 1 (**I**) signals as in **C**. Results are mean ± SD, n=3 independent experiments. **J.** Vero E6 cells transfected for 72 h with the indicated siRNA (siCtrl or combine siZDHHC8/9 and 20 - siMixT) and infected with SARS-CoV-2, 24 h. MOI≈0.1. Samples were processed as **CD** and analyzed by western blot against Spike, ERGIC53 and Cav1. **K**.**L.M** Quantification of Spike (all forms) (**K and L-**representative western blots in Figure S4J) and ERGIC53 (**M**) in each fraction as in **C**. Results are mean ± SD, n = 3 independent experiments. All *p* values were obtained by two-way ANOVA with Sidak’s multiple comparison test except for **E** where one-way ANOVA with tukey’s multiple comparison test was used.

### S-acylation of Spike drives its association with ordered lipid nanodomains

Given the abundance of Spike trimers in viruses, and the predicted compaction of the saturated acyl chains around the TMD, the local lipid organization of Spike is expected to be ordered. Consistently, it was previously observed that transfected Spike associates with detergent resistant membranes (DRMs), as we could confirm (Figure S4BC). DRMs, of which caveolin-1 is a marker, have been considered a useful biochemical readout for the association of proteins with cholesterol-rich lipid domains, even though not equivalent to the ordered lipid domains in cells (Levental et al., 2020). Such domains are predominantly found in the plasma membrane, the endosomal system and the late Golgi (van Meer et al., 2008). The ER and the early Golgi are presumed to be devoid of such domains since their cholesterol and sphingolipid contents are very low. Therefore the ERGIC, the intermediate compartment between the ER and the Golgi, into which CoVs bud (Klein et al., 2020), is not expected to contain cholesterol-rich lipid domains.

To probe the lipid environment of Spike, we first tested whether transfected intracellular Spike could be found in DRMs. For this, we biotinylated the surface proteins of transfected cells, prepared DRMs and performed streptavidin pull downs on each fraction to separate plasma membrane (bound) from intracellular (unbound) proteins. Remarkably, intracellular WT Spike was exclusively found in the DRM fractions, while cell surface WT Spike was distributed throughout the gradient (Figure 4CD). The intracellular 10C-A mutant in contrast was detergent-soluble (Figure S4DE) and also less expressed at the cell surface (Figure S4FG).

We next analyzed the distribution of cholesterol in Spike-expressing cells using the cholesterol-binding fungal metabolite, filipin. Filipin staining was more than 2-fold higher in WT than in acylation-deficient 10C-A Spike-expressing cells (Figure 4EF). In addition, staining was atypically concentrated in the perinuclear area (Figure 4F). Note that as previously observed, expression of WT, but not 10C-A Spike, promoted cell-to-cell fusion leading to the formation of large multi-nucleated syncytia (McBride and Machamer, 2010) (Figure 4F and S4I).

We next performed the DRM analysis on SARS-CoV-2 infected cells. Spike was then exclusively found in DRMs (Figure 4GH), without needing to separate intracellular from surface expressed proteins. Spike-DRM association appeared mostly intracellular as indicated by its resistance to cholesterol extraction from the plasma membrane using extracellular addition of ß-methylcyclodextrin (ßMCD). Partitioning of the viral M-protein also remain unaltered, while ßMCD treatment did solubilize caveolin-1 which is predominantly found at the plasma membrane (Figure 4G-I). The DRM association of Spike was lost when cells were treated with the MixT siRNA pool or ZDHHC20 siRNA (Figure 4J-L and S4I). In contrast, silencing of ZDHHC9 had no effect (Figure 4L and S4I). Thus ZDHHC20-mediated acylation of Spike in SARS-CoV-2 infected cells leads to it association with intracellular DRMs, presumably derived from the ERGIC, as well as from the newly formed virions on their way to the extracellular space (Figure 3H).

As mentioned, the ERGIC is not expected to have cholesterol and sphingolipid rich ordered domains. The presence of acylated Spike in DRMs in infected cells therefore raised the possibility that viral proteins drive the formation of lipid domains in the ERGIC. To test this hypothesis, we monitored the behavior of ERGIC53, a transmembrane protein that accumulates in the ERGIC. In uninfected cells, ERGIC53 was detergent soluble (Figure S4J) as expected. In contrast, ERGIC53 was mostly detected in DRMs when cells were infected with SARS-CoV-2 (Figure 4JM). This was due to protein acylation since ERGIC53 was detergent soluble in infected cells treated with the siRNA pool MixT, or siZDHHC20 but not siZDHHC9 (Figure 4JM and S4I).

Altogether this analysis suggests that ZDHHC20-mediated acylation of Spike in infected cells drives the formation of ordered domains in the ERGIC.

### S-acylation of Spike controls the local lipid environment in viral-like particle

We next investigated whether S-acylation of Spike influences its lipid environment in viral particles. We generated and purified Spike-pseudotyped viral-like particles (VLPs) from HEK293T cells, using an HIV-derived lentivector-based system. First, we verified that Spike present in VLPs is acylated (Figure S5A). Depletion of individual ZDHHC enzymes (8, 9 and 20) or all three combined (siMixT) did not significantly affect VLP production (measured by ELISA against the HIV antigen p24), incorporation of Spike into them, nor the ratio between cleaved S2 fragment and full-length Spike (Figure S5B-E). ZDHHC silencing however drasticall affected the ability of Spike to co-fractionate with DRMs (Figure 5A-C). More specifically, DRM association of Spike was lost when silencing ZDHHC20, reduced when silencing ZDHHC8, and unaltered when silencing ZDHHC9 (Figure 5C and Figure S5F).

**FIGURE 5:**
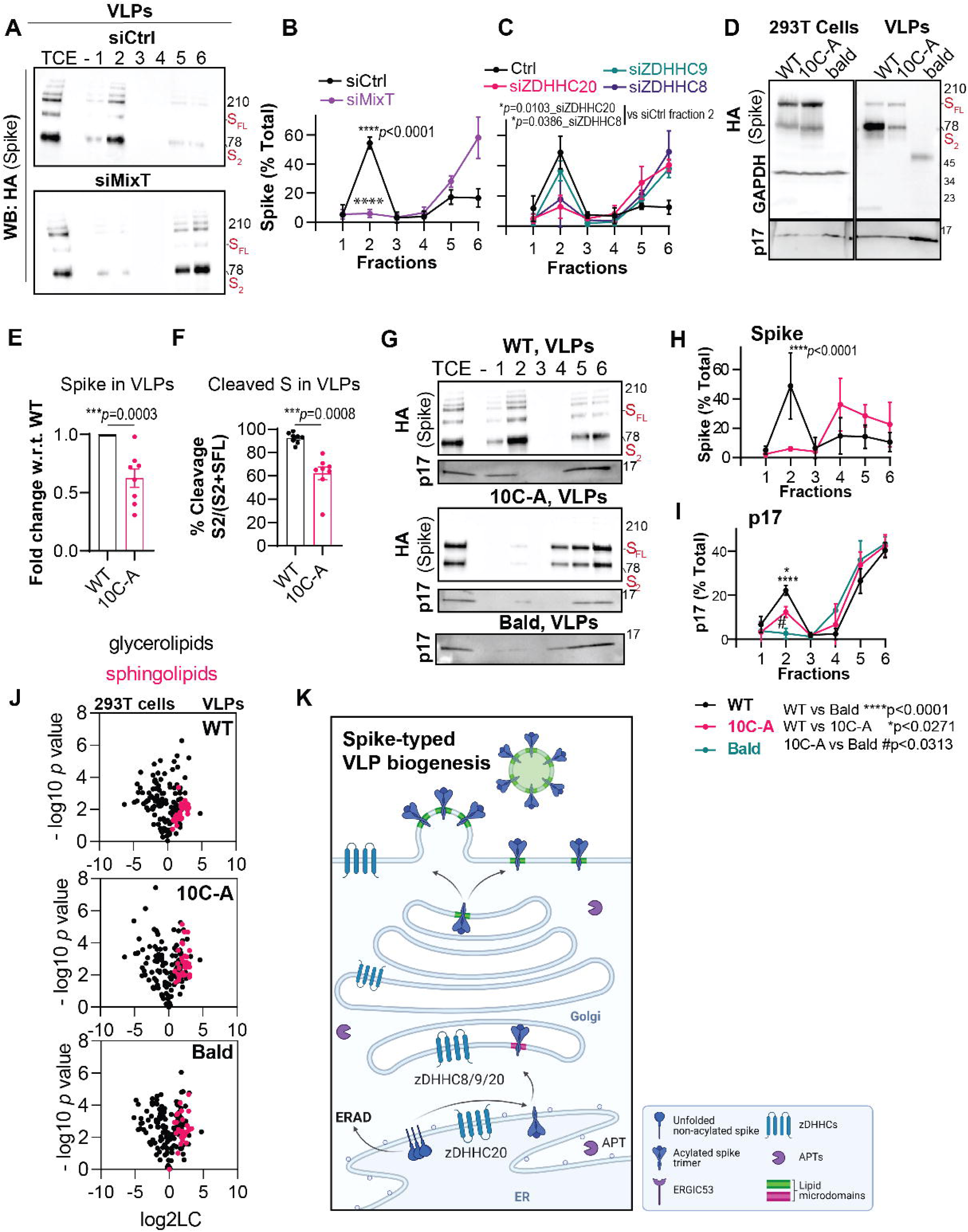
Spike S-acylation controls lipid organization of VLPs. **See alos Figure S5. A-C.** HA-Spike pseudotyped VLPs were purified from HEK293T cells depleted with the indicated siRNA (siCtrl, single or combine siZDHHC8/9 and 20 - siMixT). Concentrated VLPs were processed for fractionation and DRM isolation (see experimental procedures) and fractions analyzed by western blot for HA (Spike). **B, C.** Quantification of Spike-HA (all forms) in each fraction (obtained in **A** and **Figure S5F**) divided by the total signal of Spike-HA in the 6 fractions. Results are mean ± SD, n=3 independent experiments. **D, E, F.** Western blot analysis, and quantification of total cell lysates and correspondent VLPs: non-typed (bald) or VLPs pseudotyped with HA-Spike (WT or 10C-A mutant). Blots were analyzed for HA Spike, GAPDH (cellular-control), and p17 lentiviral matrix protein (VLP-control). **E.** HA-Spike levels were normalized against p17 levels and set to 1 for WT. **F** Spike cleavage was quantified as the ratio between S2 and S2+S full length. Results are mean ± SEM, and each dot represents an independent VLP preparation of n=8. **G, H, I.** Western blot analysis and correspondent quantification (**I**-Spike and **J**-p17) of HA-Spike (WT or 10C-A mutant) pseudotyped VLPs purified and processed for fractionation as in **A.** Results are mean ± SD, n=3 independent experiments. All *p* values were obtained by two-way ANOVA with Sidak’s multiple comparison test **(B, C, H, I),** or students t test **(E, F)**. **J.** Volcano plots of the lipid molecular species specifically enriched in VLPs, Spike-pseudotyped (WT or 10C-A mutant) or non-typed (bald), vs the correspondent producer cells (293T Cells). The log2 fold change (FC) of 293T cells vs. VLPs is plotted against the -log10 *p*-value. Black dots represent gycerolipid lipid species comparably distributed between cells and VLPs whereas red dots depict sphingolipid species, specifically enriched in VLPs. **K.** Model of S-acylation-dependent control of membrane lipid organization by Spike during cellular VLP biogenesis. Created with BioRender.com

We next generated VLPs containing either WT or 10C-A Spike, or made non-typed (bald) viruses (Figure S5I). Both forms of Spike could be efficiently detected in HEK293T cells, with a slight increase for the 10C-A mutant (Figure 5D and S5G), and displayed an equivalent ratio of cleaved vs. full length protein (Figure S5H). In contrast, Spike incorporation (Figure 5EF) and the ratio of cleaved vs. full length (Figure 5EG) was reduced by more than 30% in 10C-A containing VLPs. Such a decrease in Spike incorporation was not observed when ZDHHC expression was modified using siRNAs, likely because the silencing is incomplete and as observed acylation is highly efficient. As in MixT-treated cells, 10C-A Spike did not partition into DRMs within VLPs (Figure 5GH). Interestingly, the HIV p17 matrix protein also associated with DRMs when acylated Spike was present (Figure 5GI). Thus, as observed with ERGIC53, acylation of Spike influences its lipid environment in VLPs.

To test whether this is a local effect around Spike or a broader effect on the lipid composition of the entire VLP membrane, we performed liquid chromatography-mass spectrometry (LC-MS)-based shotgun lipidomics of purified VLPs and their producing HEK293T cells. Comparison of the relative lipid levels indicated an enrichment of sphingolipids in the VLPs, irrespective of the presence or S-acylation of Spike (Figure 5J). Thus, S-acylation of Spike induced localized formation of ordered lipid microdomains restricted to the local environment of Spike in VLPs (Figure 5K).

### Early Golgi sphingolipid composition of SARS-CoV-2 virions

We next extended our biochemical analyses to SARS-CoV-2 virions. Virions were purified from infected Vero E6 cells (48 h post inoculation) silenced for acyltransferase expression using the MixT pool. Similar levels of Spike and M in virions and cells were observed irrespective of acylation, with a small, yet, significant decrease of the Spike to M ratio in siMixT-derived virions (Figure 6AB). The relative proportion of the S2 fragment compared to the total amount of Spike was also largely unaltered by siMixT (Figure 6C). Well defined viruses were observed by cryo-electron microscopy under both conditions, with clearly visible Spikes complexes in the characteristic prefusion conformation (Figure 6D). The virions obtained from siMixT-treated cells, however, appeared less homogenous in size and slightly but significantly larger on average (Figure 6DE). We modelled the tails of the distribution of virion diameters using a Generalized Pareto Distribution, which enables the analysis of the exceedances of a particular dataset. The shape factor **(ξ)** for virions derived from control cells was negative, of −0.20 ±0.19, indicating a low propensity to generate extreme diameters (Figure 6E and S2A). In contrast, for virions derived from siMixT-treated cells, **ξ**was 0.64 ±0.34, clearly showing an increased frequency of extreme diameters (Figure 6E and S2B). Thus, ZDHHC9 and 20 contribute to the tight control of viral particle dimensions.

**FIGURE 6:**
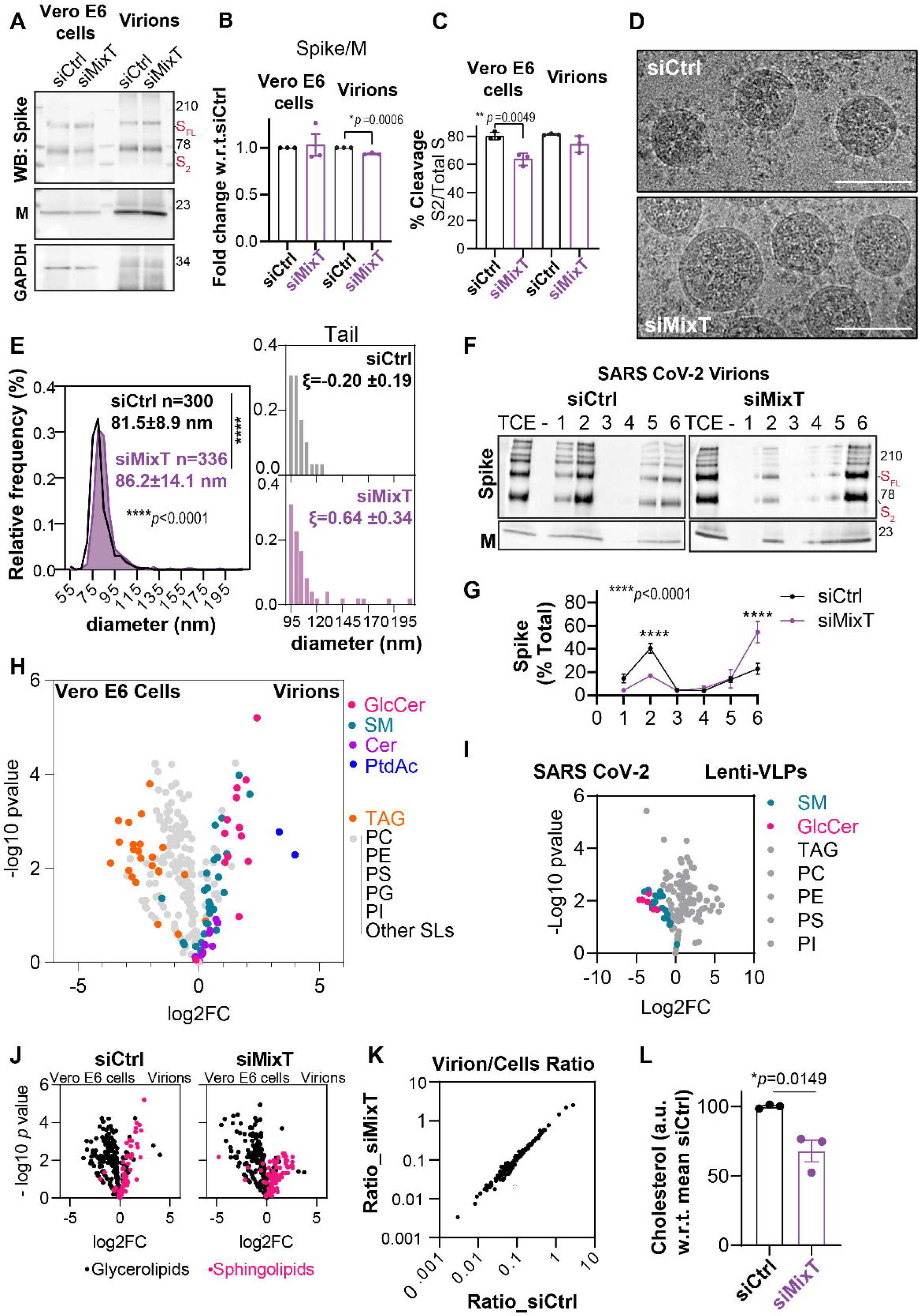
Impact of S-acylation on SARS-CoV-2 viral biogenesis. **See alos Figure S6.** For **A to I** SARS-CoV-2 virion suspensions were, purified and concentrated from Vero E6 cells transfected for 72 h with control (siCtrl) or combine siRNA to silence ZDHHC8,9 and 20 A (siMixT) and infected for 48 h with SARS-CoV-2 virus, MOI≈0.1. **A**. Western blot analysis of purified concentrated virions and total cell extracts from correspondent producer cells. Blots were probed for Spike, M and GAPDH. **B, C.** Quantification of the Spike/M ratio, set up to 1 for Controls, and the ratio between S2 and S2+S full length in Vero E6 infected cells and in Virions obtained in **A**. Results are mean ± SEM, and each dot represents an independent preparation of n=3. **D.** Virion suspensions were fixed in PFA-4%, concentrated by ultracentrifugation and analyzed by Cryo-electronic Microscopy. Scale bars= 100 nm. **E.** Frequency distribution of virion diameters (Bin width 5 nm), quantified over 300 (siCtrl) and 336 virions (siMixT) harvested from 2 independent preparations. Results are mean ± SD. A Generalized Pareto Distribution analysis of the tail frequency shows an increase of extreme diameters (shape factor ξ) for siMixT samples. **F.** Concentrated virions were processed for fractionation and DRM isolation, and fractions analyzed by western blot against Spike and M **G**. Quantification of Spike (all forms) in each fraction obtained in **F** divided by the total Spike signal in all 6 fractions. Results are mean ± SD from n=3 independent experiments. **H.** Volcano plots of lipid molecular species levels in producer Vero E6 cells vs Control SARS-CoV-2 virions. The log2 fold change (FC) is plotted against the -log10 *p*-value. Dots represent lipid species specifically enriched in virions (GlucosilCeramide, GlcCer, pink; Sphingomyelin, SM, green; Ceramides, Cer, purple; and Phosphatic Acid, PtdAc, Blue). Whereas Triglycerides, TAG, orange, are enriched in cells. In grey; PC-phosphatidylcholine; PE-phosphatidyletanolamine; PS-phosphatidylserine; PG-phosphatidylglycerol; PI-phosphatidylinositol and other SL-sphingolipids **I.** Volcano plots of lipid molecular species as in H for SARS-CoV-2 virions vs 293T-derived Spike pseudotyped VLPs. Red and green dots depict enrichment of GlcCer and SM species in SARS-CoV-2 virions. **J.** Volcano plots as in H (Vero E6 cells vs virions) depicting virion-specific sphingolipid enrichments (red dots) **(K)** and relative virion/cell lipid ratios comparisons for SARS-CoV-2 virions produced from siCtrl or siMixT silenced Vero E6 cells. **L.** Fluorometric cholesterol and cholesterol-ester analysis of lysates from SARS-CoV-2 virions produced from siCtrl or siMixT silenced Vero E6 cells. Equal volumes were used for analysis depicting comparable levels of M protein as in **A**. Results are mean ± SEM, and each dot represents an independent preparation of n=3. All *p* values were obtained by **(B, C, E, I)** students t-test and **(G)** two-way ANOVA with Sidak’s multiple comparison test.

We next analyzed the DRM association of Spike within SARS-CoV-2 envelopes. Virions were submitted to detergent solubilization in the cold. Note that for both VLPs and virions, the detergent to membrane ratio is higher than when solubilizing cells, due to the low amount of material, which generates a high risk of over-solubilization. Nevertheless, Spike was co-fractionated in DRMs in a MixT dependent manner, as observed for VLPs (Figure 6FG).

We next determined the lipid composition of SARS-CoV-2 virions and compared it to that of parental cells. We found that virions were deprived of non-membrane lipids (triglycerides >80% less represented), poor in glycerolipids (25% less represented), and enriched in sphingolipids (40% more represented) (Figure 6H and S6C-F). Virions were also enriched in phosphatidic acid which displays a small head-group and a bulky acyl tail that contributes to negative membrane curvature and may facilitate intraluminal vesicle budding (Egea-Jimenez and Zimmermann, 2018; Zhukovsky et al., 2019). More strikingly, hexosylceramide (HexCer) species (most likely glucosylceramide-GlcCer) were found to be on average 330% more represented in virions than in parental cells (Figure 6H and S6C). GlcCer, the precursor of glycosphingolipids, is produced on the cytosolic side of early Golgi *cisternae* where GlcCer synthase resides (Halter et al., 2007). Once produced, GlcCer is translocated to the lumenal leaflet of either Golgi *cisternae* or, following non-vesicular transport, of the TGN (D’Angelo et al., 2013, 2007). In the luminal leaflet of the Golgi/TGN membrane, GlcCer is readily converted to lactosylceramide and complex glycosphingolipids. Thus, GlcCer primarily populates the early segment of the biosynthetic pathway. It is widely accepted that virions retain/select a lipid composition for their envelope that is characteristic of their budding site, and this has been used as a mean to determine the lipid composition of the plasma membrane of polarized cells for example (Heaton and Randall, 2011; Ketter and Randall, 2019). Our analysis therefore provides the first detailed insight into the lipid composition of the SARS-CoV-2 budding site and shows biochemically that these viruses bud in the early secretory pathway, consistent with morphological studies (Klein et al., 2020; Stertz et al., 2007). Although the preparations of VLPs and SARS-CoV-2 virions were generated from different cells, HEK293 and Vero E6 respectively, we did attempt comparing their lipid composition. While this comparison is to be taken *cum grano salis* (with a grain of salt), it indicates that SARS-CoV-2 virions are enriched both in sphingolipids and in GlcCer when compared to VLPs. Consistent with findings on VLPs, we found that the overall lipid composition of the virions is not markedly affected by protein acylation (Figure 6JK and S6C-F). We also monitored the relative cholesterol amounts in our viral preparations normalized to the M protein. Our analysis revealed that virions derived from siMixT-treated cells contained approximately 30% less cholesterol than control virions (Figure 6L), fully consistent with the prediction that Spike S-acylation promotes the formation of ordered lipid nanodomains in the ERGIC (Figure 4).

Thus, S-acylation does not drastically affect the overall lipid composition of SARS-CoV-2 viruses but instead modifies the lipid environment in the vicinity of Spike promoting the formation of cholesterol-rich lipid domains (model in Figure 3H). This local effect is consistent with the fact that Spike trimers are rather sparsely distributed within the viral membrane as shown by cryo-tomography studies (Ke et al., 2020; Klein et al., 2020; Yao et al., 2020).

### S-acylation is essential for SARS-CoV-2 infectivity

Having characterized the biochemical effects of S-acylation on the lipid composition and organization in VLPs and SARS-CoV-2 virions, we set up to address how S-acylation affects infection. We used three complimentary approaches. First, to focus specifically on the role of Spike, we used the Spike-pseudotyped VLP system and monitored the receptor binding and fusion/entry steps. Secondly, we studied the impact of S-acylation on full SARS-CoV-2 virus by: on one hand, monitoring how wild-type viruses can infect cells depleted of specific ZDHHC enzymes; on the other, how viruses derived from siZDHHC-silenced cells can infect naïve control cells, the latter addressing the infectivity of virus produced with altered protein acylation.

To analyze binding to the ACE-2 receptor, we compared the competitive binding between Spike pseudotyped VLPs (WT or 10C-A) and purified Spike receptor binding domain (RBD), using a modified human ACE2 ELISA kits. Both WT and 10-CA pseudotyped VLPs displayed equivalent levels of competition with purified RBD, whereas bald VLPs did not display significant binding (Figure 7A). Next, we performed synchronized fusion assays by incubating ACE-2-expressing HeLa cells with VLP supernatants at 4°C, washing unbound particles and allowing subsequent fusion at 37°C for 30 min. Prior to cell harvesting, surface-attached particles were removed using a stringent acid-wash and fused and/or internalized particles were monitored by western blot. While it was possible to detect ACE2-dependent fusion of WT Spike-S2, no Spike 10C-A was detected, indicating that 10C-A-pseudotyped viral particles were unable to fuse with target cells (Figure 7B and S7A). We also monitor the infectivity of these VLPs by transducing them into ACE2-expressing HEK293 cells and monitoring luciferase expression after 3 days. The infectivity of WT Spike-pseudotyped VLPs was more than 4 times higher than that of the same amount of 10C-A containing VLPs, or bald, non-typed particles (Figure 7C). In line with this, Spike-pseudotyped VLPs, but not VSVG-pseudotyped VLPs (Figure S7B), produced from cells silenced for ZDHHC20 also displayed lower infectivity when compared to VLPs obtained from control cells (Figure 7D). Silencing ZDHHC8 or ZDHHC9 did not significantly compromise the infectivity of VLPs. Thus, ZDHHC20-mediated acylation of Spike drives its association with lipid domains, promotes its incorporation into VLPs, and increases fusion and infectivity of the VLPs.

**FIGURE 7:**
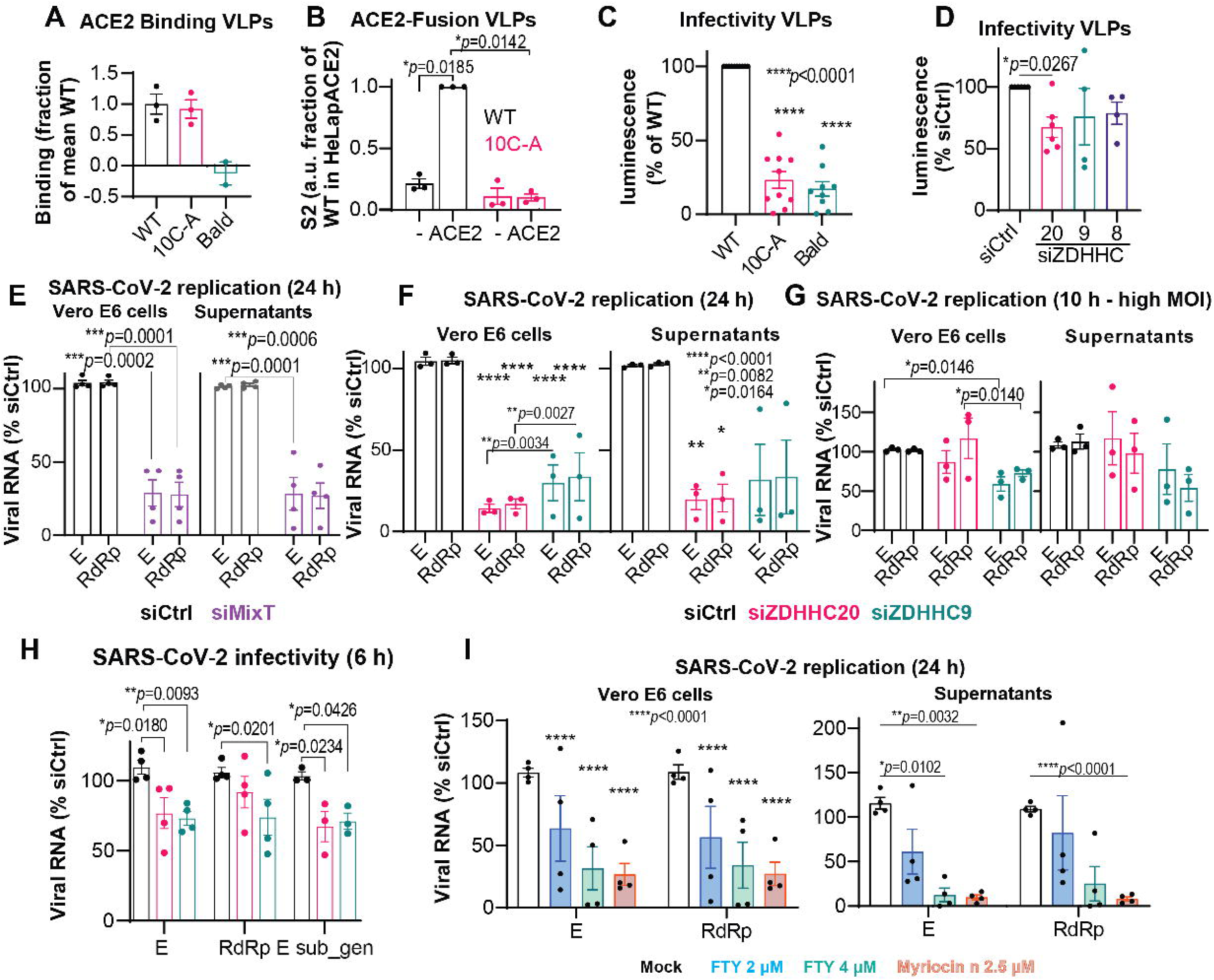
S-acylation enhances for Spike-typed VLP and SARS-CoV-2 infectivity. **See alos Figure S7. A-D.** Non-typed (bald) or HA-Spike-pseudotyped (WT or 10C-A mutant) VLP suspensions were harvested from standard HEK293T or cells transfected (72 h) with Control (siCtrl) or ZDHHC8,9 or 20-targeting siRNAs. **A**. Suspensions, with comparable Spike and p17 concentrations **(see Figure S7A)**, were used for ACE-2-based ELISA competitive-interaction assays against purified HRP-tagged Spike-receptor binding domain (RBD) (see experimental procedures). Quantitative inhibition of RBD binding was compared to culture medium and normalized to the mean of WT-pseudotyped VLP values. **B.** VLP preparations were also used for synchronized fusion assays (see experimental procedures) using HeLa or ACE-2-HeLa cells. Surface attached particles were removed by acid wash and internalized Spike S2 signal quantified by western blot **(see Figure S7A)** and expressed as fraction of WT-pseudotyped VLP signal in HeLapACE-2 cells. **AB.** Results are mean ± SEM, and each dot represents an independent preparation, n=3. **C, D.** VLPs suspensions with adjusted p24 content were used to transduce 293T-ACE2. Transduction efficiency (Infectivity) was quantified 3 days post inoculation, by lumiscence based assays. Results, normalize to WT or Ctrl conditions are mean ± SEM, and each dot derived from an independent assay of n=7 for **C** and n≥4. **E, F, G** Vero E6 transfected (72 h) with Control (siCtrl) or individual (ZDHHC9 or 20) or combine ZDHHC8,9 and 20 (siMixT) siRNA oligos were infected with SARS-CoV-2 MOI≈0.1 for 24 h or. Cells and correspondent supernatants were harvested, processed for RNA extraction and Viral RNA, for E and RdRp, quantified by q-PCR. Results are mean ± SEM and each dot represent the average RNA levels of an individual independent experiment quantified for three biological replicates for each sample. **G.H.** Vero E6 cells transfected as in **F** were infected with high SARS-CoV-2 using a MOI≈ (1-1.5) for 10 h. At 10 h post inoculation cells and correspondent supernatants were harvested for quantification of Viral RNA as in **(G, H)** and Viral proteins Spike and M **(see Figure S5C).** Virion supernatants from Control and ZDHHC9 or 20 silenced cells with comparable viral protein content and equivalent number of E copies per ml **(see Figure S5D)** were used to re-infect standard Vero E6 cells using 50 viral E copies per cell. Viral infectivity was assessed by quantifying total viral RNA (E, RdRp) and sub genomic viral RNA (E sub gen) 6 h after inoculation. Results are mean ± SEM and each dot represent the average RNA levels of an individual independent experiment (n≥3) quantified for at least three biological replicates for each sample. **I**. Quantification of Viral RNA as in A in Vero cells treated with 2.5 µM of Myriocin for ≈5 days before and during infection or treated with the indicated concentrations of FTY720 from 1 h after viral inoculation. Control cells were treated with equivalent volume of DMSO. Results are mean ± SEM and each dot represent the average RNA levels of an individual independent experiment (n=4). *p* values were obtained by **(E)** multiple student’s t test, by **(B, F, G, H, I)** two-way ANOVA with Tukey multiple comparison test or one-way ANOVA with **(C)** Tukey’s and **(D)** Dunnets multiple comparison test **(C)**.

We next analyzed the general importance of S-acylation during infection of the SARS-CoV-2 virus itself. We first chose a 24 h infection at low MOI (≈0.1). We monitored the levels of two viral RNAs, E and RNA-dependent RNA polymerase-RdRp transcripts. Both dropped by about 75% in siMixT treated infected cells, as well as in the supernatants of these cells, which contain released virions (Figure 7E). A similar reduction in viral RNA was observed when ZDHHC20 was silenced, both in the cells and in supernatants (Figure 7F). Silencing ZDHHC9 also reduced viral RNA but to a lesser extent (Figure 7F).

These experiments indicate that ZDHHC20 and 9 strongly affect the infection cycle of SARS-CoV-2. The time point chosen (24 h) allows for 2 to 3 rounds of infection, i.e. multiple rounds of entry, replication, viral biogenesis and release. To estimate the importance of acylation for the replication and/or infectivity, we analyzed infection at a shorter time, 10 h, but this required the use of a higher MOI (≈1 to 1.5) for detection. In these conditions, no significant change in viral RNA in either cells or supernatant was detected when silencing ZDHHC20, a moderate decrease was observed in cells depleted for ZDHHC9 (Figure 7G). Consistently, the levels of Spike and M proteins in infected lysates and supernatants were also comparable between conditions (Figure S7C). Within the limitations of high MOIs and incomplete loss of ZDHHC expression following siRNA, the observation suggest that ZDHHC9 influences the first round of infection, while ZDHHC20 might not.

Next, we utilized the viral supernatants produced from the 10 h infections in cells silenced for either ZDHHC20 or ZDHHC9, and used them to infect naïve control cells. Supernatants were first adjusted to comparable amounts of E-RNA copies per ml (Figure S7D). Approximately 50 E copies were then added per target cell for infection. The infectivity was monitored 6 h after viral inoculation by measuring the total RNA of E and RdRP, as well as sub-genomic E RNA (E-sub_gen), which is a more accurate proxy for viral replication. The infectivity of virions produced from either ZDHHC20 and ZDHHC9 silenced cells was 25% to 30% lower than that of control virions, based on total and sub-genomic E RNA, respectively (Figure 7H). A similar trend was observed for RdRp RNA. These data indicate that ZDHHC20-mediated S-acylation contributes to SARS-CoV-2 infection by promoting the production of fully infectious virions.

Finally, given the here identified importance of cholesterol and sphingolipid-rich domains, we tested whether pharmacological inhibition of sphingolipid biogenesis, could affect SARS-CoV-2 infection. We used the fungal metabolite Myriocin, which targets serine palmitoyltransferase-1 at early steps of sphingosine biosynthesis and fully depletes sphingolipids in cells in 5 days. We also tested FTY720, or fingolimod, a drug approved for treatment of multiple sclerosis, which while derived from Myriocin potently affects downstream enzymes, ceramide synthases. FTY720 was found to efficiently modulate the intracellular levels of sphingolipids with shorter treatment times (Berdyshev et al., 2009; Lahiri et al., 2009). Vero E6 cells were either treated with Myriocin for 5 days and subsequently infected with SARS-CoV-2 for 24 h or first infected with SARS-CoV-2 and 1 h after inoculation exposed to FTY720 for the remaining 24 h. Both treatments, Myriocin and FTY720, phenocopied the silencing of the acyltransferases, leading to a drastic reduction of viral RNA both in the cells and in supernatants after 24 h of infection (Figure 7I). Thus, disruption of sphingolipid cascades can also be used as target to disrupt cellular infection of SARS-CoV-2.

## CONCLUSION

Membrane proteins of envelope viruses were amongst the first shown to be S-acylated, yet in-dept molecular understanding of this process and its precise consequences has lagged behind. Here, based on a combination of functional genomics, biochemical, kinetic and computational studies, we show that during SARS-CoV-2 biogenesis in infected cells, Spike protein is rapidly and efficiently S-acylated following its synthesis in the ER, primarily through the action of the ZDHHC20 enzyme. This prevents premature degradation and strongly promotes the biogenesis of Spike, which is subsequently transported to the ERGIC where it arrives with 30 acyl chains decorating each trimer. The presence of these saturated lipids drives the formation of ordered membrane domains around Spike, enriched in GlcCer, sphingomyelin and cholesterol. Using VLPs pseudotyped with WT or acylation deficient Spike, as well as SARS-CoV-2 virions produced from cells silenced for ZDHHC8/9/20 expression, we could show that acylation greatly enhances viral fusion and infectivity. Our findings are fully consistent and complementary to those by Brangwynne and co-workers who recently identified, through a screening approach, that cholesterol present in the membrane of VLPs is critical for Spike-mediated fusion (Sanders et al., 2020). Our study shows that S-acylation, and the ZDHHC20 enzyme in particular, constitutes a promising drug target for coronavirus infection. Interestingly, ZDHHC20 is also involved in the acylation of the hemagglutinin of Influenza virus (Gadalla et al., 2020) and the recent elucidation of its structure indicates that this acyltransferase should be specifically druggable (Rana et al., 2018). This study sets the ground for a re-emergent interest in the study of protein S-acylation and also lipid biosynthetic pathways as important regulatory mechanisms of infection by coronaviruses and enveloped viruses in general.

## ACKNOWLEDGMENTS

We thank Machamer laboratory for M, ORF3A and E antibodies, Davide Demurtas and Graham Knott from the BioEM EPFL Core Facility for the EM analysis, Stefania Vossio and Dimitri Moreau from the ACCESS Geneva screening platform; Valeria Cagno, Caroline Tapparel and Isabella Eckerle form the University of Geneva for providing us with the SARS-CoV-2 strain and teaching us viral infections; Valerie Chavez from University of Lausanne for performing the statistical analysis of extreme values. We thank all the members of the F.G.v.d.G. lab for discussions and suggestions. We thank Panyain Nattawadee for the model design (Created with BioRender.com) and Christina Ernst for primer sharing. This work was supported by the Swiss National Science Foundation Corona Call, the CARIGEST foundation and the EPFL Corona Research task force to F.G.v.d.G. MD simulations were carried out on the Piz Daint computer at the Swiss Supercomputing Center (CSCS) thanks to access granted by PRACE Covid19 fast track project #17 (pr97) to M.D.P.

## AUTHOR CONTRIBUTIONS

Conceptualization, F.S.M., L.A., D.T., G.D’A., F.G.V.D.G.; Investigation, F.S.M., L.A., O.S., P.T., B.K., C. R., J.P.M., L.A.A., G.D’A.; Funding Acquisition, F.S.M., M.D.P. & F.G.V.D.G.; Writing–Original Draft, F.M., L.A., G.D’A., L.A.A., F.G.V.D.G.; Writing–Review & Editing, F.S.M., L.A., O.S., P.T., J.P.M., L.A.A., M.D.P., D.T., G.D’A., F.G.V.D.G.; Resources, B.K., O.S.

## STAR METHODS

### KEY RESOURSES TABLE

KEY RESOURCES TABLE
| REAGENT or RESOURCE | SOURCE | IDENTIFIER |
| --- | --- | --- |
| **Antibodies** |  |  |
| ACE2 | ABCCAM | RRID:AB_301861 Cat#ab15348 |
| ACTIN | MILLIPORE | RRID:AB_2223041 Cat#MAB1501 |
| CALNEXIN | MILLIPORE | RRID:AB_2069152 Cat#MAB3126 |
| CAVEOLIN-1 | SANTA CRUZ | RRID:AB_2072042 Cat#sc-894 |
| E | gift from Machamer lab | RRID:NA |
| ERGIC53 | ProSci | RRID:AB_796913 Cat#PSC-PM-7213 |
| FILIPIN | SIGMA | Cat#F4767 |
| FLAG | SIGMA | RRID:AB_259529 Cat#F3165 |
| GAPDH | ThermoFisher | RRID:AB_2533438 Cat#398600 |
| GIANTIN | ABCCAM | RRID:AB_880195 Cat#ab 37266 |
| GM130 | BD | RRID:AB_398141 Cat#610823 |
| HA-HRP | ROCHE | RRID:AB_390917 Cat#12013819001 |
| HA | ROCHE | RRID:AB_390918 Cat#11867423001 |
| HA gel beads | ROCHE | RRID:AB_390914 Cat#1815016001 |
| LRP6 | Cell signaling | RRID:AB_10831525 Cat#2560 |
| M | gift from Machamer lab | NA |
| MYC | SIGMA | RRID:AB_439694 Cat#M4439 |
| ORF3A | gift from Machamer lab | NA |
| Propidium iodide PI | ThermoFisher | RRID:P3566 |
| protein G al beads | GEHealthcare | Cat#17-0618-01 |
| P17 | home made, gift from Trono lab | NA |
| SPIKE | LIFESPAN | RRID:AB_840148 Cat#LS-C19510 |
| streptavidin beads | SIGMA | Cat#S1638 |
| Transferrin Receptor | ThermoFisher | RRID:AB_2533029 Cat#13-6800 |
| ZDHHC3-GODZ | ABCCAM | RRID:AB_742236 Cat#ab31837 |
| TUBULIN | SIGMA | RRID:AB_477579 Cat#T5168 |
| alexa-647-d.anti-mouse | ThermoFisher | RRID:AB_2536183 Cat#A31573 |
| alexa-488 anti-rat | Jackson ImmunoResearch | RRID:712.545.153 |
| anti-mouse HRP | GEHealthcare | RRID:AB_772210 Cat#NA931V |
| anti-rabbit-HRP | GEHealthcare | RRID:AB_772206 Cat#NA934V |
| **Bacterial and virus strains** |  |  |
| hCoV-19/Switzerland/GE9586/2020 | Switzerland-Geneve | EPI_ISL_414022\|2020-02-27 |
| **Chemicals, peptides, and recombinant proteins** |  |  |
| BREFELDIN A | BioLegend | Cat#BLG-420601 |
| CYCLOHEXIMIDE | SIGMA | Cat#C7698 |
| FTY720 | SIGMA | Cat#SML0700 |
| Hydroxylamine | SIGMA | Cat#55460 |
| beta-methyl-cyclodextrin | SIGMA | Cat#C4555 |
| MYRIOCIN | CAYMAN Chemical | Cat#TRCM884400 |
| Palmostatin B | CALBIOCHEM | Cat#178501 |
| **Critical commercial assays** |  |  |
| Amplex red cholesterol assay | ThermoFisher | Cat#A12216 |
| Amplify NAMP100 | GEHealthcare | Cat#GENAMP100 |
| AVICEL GP3515 | DUPONT | Cat#41094401 |
| Bright-Glo Luciferase assay | PROMEGA | Cat#E2610 |
| Crystal violet | SIGMA | Cat#V5265 |
| EPISERF Medium-1 | ThermoFisher | Cat#10732022 |
| DMEM | ThermoFisher | Cat#31966021 |
| DMEM w/o cystein, methionin | Bioconcept | Cat#1-26S59 |
| ECL reagent spray-HRP | Witec | Cat#K-12049-D50 |
| EZ-Link Sulfo-NHS-SS-biotin | PIERCE | Cat#PIER21326 |
| E.Z.N.A viral RNA kit | OMEGA Bio-teck | Cat#R6874-02 |
| FUGENE 6 | PROMEGA | Cat#E2692 |
| GMEM for IM | ThermoFisher | Cat#11710035 |
| HBS-EP buffer | CYTIVIA | Cat#BR100188 |
| iSCRPT | BIO-Rad | Cat#1708891 |
| Lacey carbon film EM grid | Agar Scientific UK | Cat#POL24924-25 |
| MALEIMIDE | Fluorochem | Cat#11793 |
| MEM | SIGMA | Cat#M4655 |
| MIRIUS TRANSIT-X2RTM | MIRIUS Bio Corporation | Cat#MIR6000 |
| MMTS | SIGMA | Cat#64306 |
| OPTIMEM-reduced serum medium | ThermoFisher | Cat#31985062 |
| OPTIPREP | AXISSHIELD | Cat#1114542 |
| Paraformaldehyde | Electron Microscopy Science | Cat#15714 |
| PBS | BioConcept | Cat#3-05F29-I |
| PBS-EDTA | BioConcept | Cat#5-32F00-I |
| ProLond mounting medium | ThermoFisher | Cat#P36934 RRID : SCR_015961 |
| protease inhibitor Complete | ROCHE | Cat#11836153001 |
| P24 ELISA | Zeptometrix | Cat#801200 |
| QUICKchange II XL | AGILENT | Cat#200522 |
| SARS--Cov2 surrogate virus neutralization kit | GENSCRIPT | Cat#L00847 |
| SYBR green Master Mix | Roche | Cat#4913914001 |
| TCEP | SIGMA | Cat#C4706 |
| thiopropyl sepharose beads | CreativeBiomart | Cat#Thio-001A |
| TX-100 | PIERCE | Cat#PIER28314 |
| Water Ambion | ThermoFisher | Cat#AM9906 |
| Zebra spin desalting columns | PIERCE | Cat#PIER89882 |
| 3H-palmitic acid | American Radio Labelled Chemicals | Cat#ART0129-25 |
| 35S-methionin-cysteine | Hartmann Analytic GmbH | Cat#IS103-185 |
| 5kDa PEG | SIGMA | Cat#63187 |
| **Experimental models: cell lines** |  |  |
| HEK-293T | ATCC | CVCL_1926 |
| HELA | ATCC | CVCL_0030 |
| HAP1 | Horizon discovery | CVCL_XU08 |
| VERO E6 | ATCC | CVCL_0574 |
| 293T-ACE2 | Generated in this study | NA |
| Oligonucleotides see Supplementary information |  |  |

### RESOURCE AVAILABILITY

#### Lead Contact and Material Availability

All resource/reagent requests will be available upon contact to Francisco Mesquita.

### EXPERIMENTAL MODEL AND SUBJECT DETAIL

#### Cell culture

HAP1 cells were obtained from Horizon Discovery, Vero E6 cells were obtained from the American Type Culture Collection (ATCC) and maintained in DMEM medium supplemented with 10% FCS, penicillin and streptomycin; HELA cells were maintained in MEM medium supplemented with 10% FCS and complemented with 2 mM glutamine and 1% of non-essential amino-acids. For lentiviral vector production, 293T cells were cultured in DMEM supplemented with 10% FBS for maintenance and transfection, and in EpiSerf for production. All cells were kept at 37°C in a humidified atmosphere of 5% CO_2_ and were tested negative for mycoplasma.

The 293T-ACE2 stable cell line was generated by transduction with an ACE2-ires-puro encoding lentivector (produced as described below and at a MOI of 0.5) followed by puromycin selection and maintained in DMEM 10% FCS with 1µg/ml puromycin.

#### Viral Stock production and titration with plaque-based assays

All viral stocks were produced and isolated from supernatants of Vero E6 cells, cultured in T75 culture flasks to a confluency of 80-90%, and infected with an original passage 2 (P2) SARS-Cov-2 virus, for 48 or 72 h, at MOI≈0.05, in 10 ml DMEM supplemented with 2.5% FCS. Original stocks were obtained from the following strain: hCoV-19/Switzerland/GE9586/2020|EPI_ISL_414022|2020-02-27. Passage 3 supernatants were harvested, clear of cell debris by centrifugation (500 *g* 10 min) and filtration (0.45 µm), aliquoted and stored at −80 C. Viral titers were quantified by determining the number of individual plaque forming units after 48 h of infection in confluent Vero E6 cells. In brief, viral stocks were serially diluted (10-fold) in serum-free medium and (400 μl) inoculated in triplicate 48 wells, confluent Vero E6 cells (2.5 x 10^5^) cells per well. After 1 h, inoculums were discarded and cells overlaid with a mixture of 0.4% of Avicel-3515 (Dupont) (from 2% stock) in DMEM supplemented with 5% FCS and pen/strep for additional 48 h. Overlays were discarded and cells fixed in 4% PFA for 30 min at RT. Fixed cells were washed in PBS and stained with 0.1% crystal violet solution (in 20% ethanol/water) for 15 min. Staining solution was discard and wells washed twice in water. Plates were allowed to dry and analyzed for quantification of the cytopathic effect as number of individual Plaque forming units (PFU) per ml (Avg PFU*1/Volume*1/dilution factor). Such un-concentrated viral stocks yielded between 0.5 to 3×10^6^ PFU/ml.

#### Oligonucleotides and Plasmids

HDM-IDTSpike-fixK (BEI catalog number NR-52514): plasmid expressing under a CMV promoter the Spike from SARS-CoV-2 strain Wuhan-Hu-1 (Genbank NC_045512) codon-optimized using IDT and the Kozak sequence in the plasmid fixed compared to an earlier version of this plasmid was obtained from Bloom Lab-Fred Hutchinson Cancer Research Center. Spike with mutation D614G was generated from HDM-IDTSpike-fixK using mutated oligonucleotides annealing. Plasmids expressing epitope-tagged SARS-CoV-2 Spike proteins were cloned in pcDNA6.2 with HA-tag in C-terminal using the Gateway system, and the donor plasmid pDONR223_SARS-CoV-2_Spike (Fritz Roth, Addgene plasmid # 149329; http://n2t.net/addgene:149329; RRID: Addgene_149329). Cysteine to Alanine substitutions were also generated with Quickchange.

Plasmids expressing human ZDHHC proteins were cloned in pcDNA3.1 with MYC-tag (provided by the Fukata lab).

The ACE2 sequence (from pHAGE2-EF1aInt-ACE2-WT a kind gift from J.D. Bloom) was swapped with the LacZ cassette of the pRRL-PGK-lacZ-ires-puro lentivector to generate the pRRL-ACE2-ires-puro vector.

All oligonucleotides, namely QPCR primers and siRNA oligos used in this study are described in detail in Supplementary table 1 and 2.

### EXPERIMENTAL MODEL AND SUBJECT DETAIL

#### Transfection for Plasmids and siRNA

Unless otherwise indicated, plasmids were transfected into 293T, Hela or Vero cells for 24 or 48 hours (3 µg/ 9.6cm^2^ plate) using Transit-X2RTM (Mirius) transfection reagent. For control transfections, we used an empty pcDNA6.2 plasmid.

siRNA for Hela cells or for Vero cells were purchased from Qiagen (see supplementary table). For gene silencing, cells were transfected for 72 hours with (15 pmol/ 9.6cm^2^ plate) using Transit-X2RTM (Mirius) transfection reagent. Control siRNA and species specific siRNA oligos targeting human or monkey ZDHHCs are described in Supplementary table 1.

#### SARS CoV-2 infections

All infections for experimental analysis were done using passage 3 SARS-CoV-2 stocks. Vero E6 cells seeded to a confluency of 90 to 100%, were, washed twice in warm serum-free medium and inoculated with the indicated MOI of SARS-CoV-2, diluted in serum free medium (5 ml for T75; 2 ml for T25; 1 ml for 6-well plates). 1 hour after inoculation cells were washed with complete medium and infection allowed to proceed for the indicated time points in DMEM supplemented with 2.5% FCS, penicillin and streptomycin (unless otherwise indicated: 10 ml for T75; 4 ml for T25; 2 ml for 6-well plates). Cells cultured in T75/T25 flasks for biochemical analysis, or 6-well plates for qPCR and Flow cytometry experiments, were infected with a MOI of ≈0.1, except for time-course experiments measuring: radioactive ^3^H-Palmitic acid-incorporation; - decay; and Acyl-pegylation dynamics, in which cells were infected with a MOI of ≈0.5 to 1. For infections involving ^3^H-Palmitic acid, medium without serum IM (see below) was used for all washes, viral inoculation and throughout infection.

#### Immunoprecipitation

For immunoprecipitations, cells were lysed 30 min at 4°C in IP buffer (0.5%NP40, 500 mM Tris-HCl pH 7.4, 20 mM EDTA, 10 mM NaF, 2 mM benzamidine, and a cocktail of protease inhibitors), centrifuged 3 min at 2000 g and supernatants were precleared with protein G-agarose conjugated beads and supernatants were incubated 16 h at 4°C with specified antibodies and beads.

#### Radiolabeling ^3^H-palmitic acid incorporation

To follow S-acylation, transfected or infected cells were incubated 1 hour in medium without serum IM (Glasgow minimal essential medium buffered with 10 mM Hepes, pH 7.4), followed by 2 hours or indicated hours at 37°C in IM with 200 µCI /ml ^3^H palmitic acid, washed with IM and incubated different times at 37°C with complete medium prior immunoprecipitation overnight. Beads were incubated 5 min at 90°C in reducing sample buffer prior to SDS-PAGE. Immunoprecipitates were split into two, run on 4-20% gels and analyzed either by autoradiography (^3^H-palmitate) after fixation (25% isopropanol, 65% H_2_O, 10% acetic acid), gels were incubated 30 min in enhancer Amplify NAMP100, and dried; or Western blotting. Autoradiograms and western blotting were quantified using the Typhoon Imager (Image QuantTool, GE healthcare).

#### Radiolabeling ^35^S-cys/met incorporation

For metabolic labeling, transfected cells were washed with methionine /cysteine free medium, incubated 4hours pulse at 37°C with 50 µCi/ml ^35^S-methionine/cysteine, washed and further incubated for different times at 37°C in complete medium with a 10-fold excess of non-radioactive methionine and cysteine. Proteins were immunoprecipitated and analyzed by SDS-PAGE. Autoradiography and western blotting were quantified using the Typhoon Imager (Image QuantTool, GE healthcare).

#### Drug treatments

Protein synthesis was blocked by 1 hour or different indicated time treatment with 10 µg/ml cycloheximide at 37°C in serum free medium. To block protein transport from ER to Golgi, we used Brefeldin A at 5 µg/ml for 1 hour at 37°C in serum free medium. To sequester cholesterol, beta-metyl-cyclodextrin was used at 10 mM for 30 min in serum free medium. As inhibitor of depalmitoylation, Palmostatin B was used at 10 µM for 4 hours in serum free medium. To block serine palmitoyltransferase, denovo sphingolipid biosynthesis, cells were cultured either in Myriocin at 2.5 µM, 5 days prior infection and during the course of infection, or with a synthetic myriocin analogue, Fingolimod-FTY20 at 2 µM or 4 µM during infection.

#### Acyl-RAC capture assay

Protein S-palmitoylation was assessed by the Acyl-RAC assay as previously described (Werno and Chamberlain, 2015) with some modifications. Cells or supernatants were lysed in 400 μl buffer (0.5% Triton-X100, 25 mM HEPES, 25 mM NaCl, 1 mM EDTA, pH 7.4, and protease inhibitor cocktail). Cell lysis were incubated 30 min at RT with 10mM TCEP. Then, 200 μl of blocking buffer (100 mM HEPES, 1 mM EDTA, 87.5 mM SDS, and 1.5% [v/v] methyl methanethiosulfonate (MMTS)) was added to the lysates and incubated for 4 h at 40°C to block free the SH groups with MMTS. Proteins were acetone precipitated and resuspended in buffer (100 mM HEPES, 1 mM EDTA, 35 mM SDS). For treatment with hydroxylamine (NH2OH) and capture by Thiopropyl Sepharose beads, 2 M of hydroxylamine was added together with the beads (previously activated for 15 min with water) to a final concentration of 0.5 M of hydroxylamine and 10% (w/v) beads. As a negative control, 2 M Tris was used instead of hydroxylamine. These samples were then incubated overnight at room temperature on a rotating wheel. After washes, the proteins were eluted from the beads by incubation in 40 μl SDS sample buffer with ß-mercaptoethanol for 5 min at 95°C. Finally, samples were separated by SDS-PAGE and analyzed by immunoblotting. A fraction of the cell lysate was saved as the input.

#### Acyl-Peg-exchange

This PEG switch assay is used to PEGylate previously palmitoylated cysteines following removal of hydroxylamine as previously described (Plain et al., 2020). To block free cysteine, cells or supernatant or purified virions were lysed and incubated in 400 μl buffer (2.5% SDS, 100 mM HEPES, 1 mM EDTA, 100mM Maleimide pH 7.5, and protease inhibitor cocktail) for 4 h at 40°C. To remove excess unreacted maleimide, proteins were acetone precipitated and re-suspended in buffer (100 mM HEPES, 1 mM EDTA, 1% SDS, pH 7.5). Previously palmitoylated cysteines were revealed by treatment with 250 mM hydroxylamine (NH2OH) for 1 hour at 37°C. Cell lysates were desalted using Zebra spin columns and incubated 1 hour at 37°C with 2mM 5kDa PEG: methoxypolyethylene glycol maleimide. Reaction was stopped by incubation in SDS sample buffer with ß-mercaptoethanol for 5 min at 95°C. Samples were separated by SDS-PAGE and analyzed by immunoblotting. A fraction of the cell lysate was saved before the addition of 5kDa PEG as the input.

#### Isolation of detergent-resistant membranes (DRMs)

Approximately 1 × 10^7^ cells were re-suspended in 0.5 ml cold TNE buffer (25 mMTris-HCl, pH 7.5, 150 mM NaCl, 5 mM EDTA, and 1% Triton X-100) with a tablet of protease inhibitors (Roche). Membranes were solubilized in a rotating wheel at 4 °C for 30 min. DRMs were isolated using an Optiprep^TM^ gradient: the cell lysate was adjusted to 40% Optiprep^TM^, loaded at the bottom of a TLS.55 Beckman tube, overlaid with 600 μl of 30% Optiprep^TM^ and 600 μl of TNE, and centrifuged for 2 hours at 55,000 rpm at 4 °C for cells or 4 hours at 55,000 rpm at 4 °C for VLPs. Six fractions of 400 μl were collected from top to bottom. DRMs were found in fractions 1 and 2. Equal volumes from each fraction were analyzed by SDS-PAGE and western blot analysis using anti-Spike, anti-M, HRP-conjugated anti-HA, ERGIC53, caveolin1 and transferrin receptor antibodies.

#### Surface biotinylation

Surface biotinylation was performed on transfected cells. Cells were allowed to cool down shaking at 4°C for 15 min to arrest endocytosis. Cells were then washed three times with cold PBS and treated with EZ-Link Sulfo-NHS-SS-Biotin No weight for 30 min shaking at 4°C. Cells were then washed 3 times for 5 min with 100mM NH4Cl and lysed in 1% Tx-100 to do DRMs or in IP Buffer for 1h at 4°C. Lysate were then centrifuged for 5 min at 5000rpm and the supernatant incubated with streptavidin agarose beads overnight on a wheel at 4°C. Beads were washed with IP buffer 5 times and the proteins were eluted from the beads by incubation in SDS sample buffer with ß-mercaptoethanol for 5 min at 95°C. Samples were separated by SDS-PAGE and analyzed by immunoblotting. Western blots were developed using the ECL protocol and imaged on a Fusion Solo from Vilber Lourmat. Densitometric analysis was performed using the software Bio-1D from the manufacturer.

#### Lentiviral particle production

Plasmids expressing SARS-CoV-2 Spike WT-HA or SARS-CoV-2 Spike 10C-A-HA proteins, pHAGE2-CMV-Luc-ZSgreen, Hgpm2, REV1b, Tat1b (a kind gift from J.D. Bloom) were co-transfected into 293T cells for 24 hours with the following ratio 3/9/2/2/2 (18 μg/ 56.7cm^2^ plate) using Fugene transfection reagent. The following day, cells were transferred in EpiSerf medium, and cell supernatants were collected after 8 hours and 16 hours. Harvested supernatants were pooled, clarified by low-speed centrifugation, filtered to remove cell debris (5 ml were kept at this stage for infectivity assays) and concentrated by centrifugation at 47000g through a 20%(w/v) sucrose cushion for 90min. Concentrated virus like particles (VLPs) were immediately resuspended in 300 μl of cold TNE buffer (25mM Tris-HCl, pH 7.5, 150 mM NaCl, 5 mM EDTA, and 1% Triton X-100) for biochemical experiments.

#### Binding Assay ELISA

Binding between Spike-pseudotyped VLPs and ACE-2 was determining using an ELISA-based SARS-CoV-2 Surrogate Virus Neutralization Test Kit with a modified protocol. In brief, Non-concentrated VLPs suspensions adjusted to comparable Spike and p17 content (monitor by western blot) were used to pre-incubate human-ACE2-coated ELISA plates for 30 min at 37°C. After 30 min, 100 μl of diluted (1/2000) Horseradish peroxidase (HRP)-conjugated Spike Receptor binding domain RBD was added to each well for further incubation, 15 min at 37°C. Plates were then washed 4x in the Kits washing solution and developed with 100 μl of substrate solution for 10 min in the dark, followed by a quenching step with 50 μl of stop solution for each well. Absorbance was monitored at 450 nm immediately and inversely correlated with the binding of Spike VLPs to ACE-2. Eppiserf medium was used as negative control, and results expressed as % binding = (1 - (Sample OD/ Neg.Control OD)) normalized to 1 for the mean of WT-Spike-pseudotyped VLPs.

#### Synchronized fusion assay

ACE-expressing HeLa or standard HeLa cells seeded in six well plates were washed 2 times in cold PBS and incubated on ice in cold Eppiserf medium for 2 to 5 min. Cells were then incubated with equivalent volumes of non-concentrated VLPs suspensions adjusted to comparable Spike content (monitored by western blot) for 30 min on ice to allow binding. Cells were subsequently washed 3 times in cold PBS and incubated in warm Eppiserf medium at 37°C for 30 min to allow fusion. Next, cells were washed 2 times in cold PBS and incubated in Acid-wash buffer (145mM NaCl, 20mM MES-Tris pH 4.5) for 10 min with gentle rocking, to remove unfused surface attached particles. Cells were then lysed 30 min at 4°C in IP buffer, centrifuged 3 min at 2000 g and supernatants resuspended in SDS sample buffer with ß-mercaptoethanol for 5 min at 95°C. Samples were separated by SDS-PAGE and analyzed by Western blot.

#### VLPs Infectivity

Non-concentrated VLPs suspensions were adjusted for p24 content (as monitored by ELISA) and used to transduce 293T-ACE2 cells (technical replicates n=12). Transduced cells were analyzed 3 days later with the Bright-Glo Luciferase assay system.

#### Analysis of intracellular viral RNA-Q-PCR

For time-course optimization (Fig S2) and 24 h replication experiments (Fig 7) Infected Vero-E6 cells and correspondent supernatants (150 μl), cultured in 6-well plates (2 ml culture medium per well - 3 independent wells per condition) were harvested at indicated time points and lysed in 500 μl of mastermix QVL lysis buffer from E.Z.N.A. Viral RNA kit used for Viral RNA extraction according to manufacturer’s instructions. RNA concentration was measured and 500 ng or 1000 ng of total RNA was used for cDNA synthesis using iScript. A 1:5 dilution of cDNA was used to perform quantitative real-time PCR (QPCR) using Applied Biosystems SYBR Green Master Mix on 7900 HT Fast QPCR System (Applied Biosystems) with SDS 2.4 Software. Primers used are described in Supplementary table 2. All data (always in triplicate) were normalized to Ct values from three housekeeping (HK) genes ALAS-1, Guss and TBP, except for supernatants from replication experiments at 24 h, which Ct data were normalized using Ct values from correspondent cellular HK genes. Results were expressed as 2^(-ΔΔCt)*100%.

#### Titration of Viral E copies

For titration of viral RNA, equivalent volumes of RNA extracted from infected culture supernatants or RNA from serial dilutions of in vitro transcribed RNA for Wuhan coronavirus (2019-nCoV) targeting region of the E gene standards (E-standards - European Virus Archive platform - Ref. 026N-03866) were used for cDNA synthesis. Samples were then used to perform QPCR analysis (using E specific primers) as previously described. A standard curve was generated by plotting the Ct values against the number of E copies per ml indicated by the serial dilution of the E-standards (stock at 1e5 copies / µl) and used to extrapolate the number of E copies per ml in samples.

#### SARS-CoV-2 single round infections and infectivity analysis

Vero-E6 cell cultured in T75 culture flasks (≈80-90% confluency after 72 h siRNA transfection) were infected with MOI=1 as described and further incubated in 5 ml culture medium per flask using 3 independent flasks per condition. 10 h after inoculation cells were washed scrapped and harvested with correspondent supernatants. Cells (1/2 of T75) and correspondent supernatants (150 μl) were processed for QPCR analysis or lysed in IP buffer and processed for Western blot as described. Remaining supernatants were aliquoted and stored at −80°C for titration of viral E copies. After titration and adjustment to viral E RNA copies non-concentrated SARS-CoV-2 supernatants were used to infect confluent Vero E6 cell monolayers cultured in 12 well plates using an approximate ration or 50 E copies per cultured host cell. At least 3 wells were infected per supernatant per condition. Infection was done as described until 6 h post inoculation when cells were washed lysed and processed for QPCR analysis as described

#### Analytical flow cytometry of SARS-CoV-2 infected cells

Vero E6 cells were infected in 6-well plates [as mentioned previously]. Cells washed and scrapped in 1 ml PBS and recovered by mild centrifugation 400g 3 min. Supernatants were discard and cells were fixed in 4% PFA for at least 30 min, washed twice in PBS, and permeabilized for 5 min with 0.1% Triton in PBS. The following blocking, antibody, and washing steps were all done with FACS buffer (2% FBS in PBS/EDTA) primarily at RT. Blocking was done for 15 min, primary antibody incubation with mouse anti-Spike antibody (LC-C19510) was performed at 1:200 for 30 min at RT or at 1:500 ON at 4 °C, and cells were washed twice. Alexa-647-conjugated donkey anti-mouse was used as a secondary antibody at 1:600 for 30 min at RT, after which cells were washed three times and kept cold before flow cytometry acquisition. Analytical flow cytometry was performed using an LSRII or LSR Fortessa (BD; Becton Dickinson) instrument and results were analyzed using the FlowCore package in R.

#### Immunohistochemistry

Vero E6 cells seeded in glass coverslips in 24-well plates and transfected with the indicated myc-ZDHHC or HA-Spike constructs (for 24 h) were fixed in 4% paraformaldehyde (15 min), quenched with 50 mM NH4Cl (30 min) and permeabilized with 0.1% Triton X-100 (5 min). Antibodies were diluted in PBS containing 1% BSA. Coverslips blocked in PBS 1% BSA, for 30 min, washed and incubated with primary antibodies for up to 2 hours RT or overnight at 4°C, washed three times in PBS, and incubated 45 min with secondary antibodies. Coverslips were mounted onto microscope slides with ProLong™ Gold Antifade Mountant. Images were collected using a confocal laser-scanning microscope (Zeiss LSM 700) and processed using Fiji™ software.

#### SARS CoV-2 isolation for biochemical, LPDX, Cryo-EM, Infectivity assays

Vero E6 cells cultured in T75 flasks, transfected with the indicated siRNAs were infected as described. 1 h after viral inoculation cells were washed twice and incubated in EpiSerf medium (serum free) for the remaining of the infection. Culture supernatants (10 ml per flask) were harvested after 48 h, clarified by low-speed centrifugation (500 g 10 min) and filtration (0.45 µm); and stored at −80 C (for infectivity assays), or further processed for biochemical, LPDX or CryO EM experiments.

#### Electronic Microscopy

At least 20 ml (10 ml per T75 flask per condition per preparation) of pre-cleared culture supernatants from 48 h infections were inactivated with paraformaldehyde (PFA; final concentration 4%) for > 2 h and concentrated by centrifugation at 47000g through a 20%(w/v) sucrose cushion for 90min 16°C. Concentrated virions were immediately re-suspended in cold 50 µl of EM-gradel Hepes buffered saline for subsequent EM processing. A Lacey carbon film EM grid was held in tweezers and 4-5 μL of sample solution was applied on the grid. The tweezers are mounted in an automatic plunge freezing apparatus (Vitrobot, ThermoFisher) to control humidity and temperature. After blotting, the grid was immersed in a small metal container with liquid ethane that is cooled from outside by liquid nitrogen. The speed of cooling is such that ice crystals do not have time to form. Observation was made at −170°C in a Tecnai F 20 microscope (ThermoFisher) operating at 200 kV equipped with a cryo-specimen holder Gatan 626 (AMETEK). Digital images were recorded with a Falcon III (ThermoFisher) camera 4098 × 4098 pixels. Magnification of 50000X with a pixel size of 0.2nm, using a defocus range from −2µm to −3µm.

#### Concentration of SARS-CoV-2 virions for LPDX and biochemical assays

Approximately 10 ml (per T75 flask) of pre-cleared culture supernatants from 48 h infections were aliquoted into 1.5 ml microtubes and concentrated by centrifugation at 47000g through a 20%(w/v) sucrose cushion (200 μl) for 90min. Medium and sucrose supernatants were discard, pellets allowed to dry and re-suspended and pooled in: 400 μl of cold 1%-Triton X TNE buffer for DRM analysis; or 500 µl 1:1 mixture of methanol:chloroform for LPDX analysis.

#### Lipid extraction

Total lipid extracts were prepared using a standard MTBE protocol followed by a methylamine treatment for total lipid analysis by mass spectrometry. Briefly, cell pellets or viral fractions were resuspended in 100 μL H2O. 360 μL methanol and 1.2 mL of MTBE were added and samples were placed for 10 min on a vortex at 4°C followed by incubation for 1 h at room temperature on a shaker. Phase separation was induced by addition of 200 μL of H2O. After 10 min at room temperature, samples were centrifuged at 1000 g for 10 min. The upper (organic) phase was transferred into a glass tube and the lower phase was re-extracted with 400 μL artificial upper phase [MTBE/methanol/H2O (10:3:1.5, v/v/v)]. The combined organic phases were dried in a vacuum concentrator. Lipids where then resuspended in 500 μL of ChCl3.

#### LC-MS untargeted lipidomics

Lipid extracts (2 μL injection volume in ChCl3:MeOH 2:1) were separated over an 8 min gradient at a flow rate of 200 μL/min on a HILIC Kinetex Column (2.6lm, 2.1 × 50 mm2) on a Shimadzu Prominence UFPLC xr system (Tokyo, Japan). Mobile phase A was acetonitrile:methanol 10:1 (v/v) containing 10 mM ammonium formate and 0.5% formic acid while mobile phase B was deionized water containing 10 mM ammonium formate and 0.5% formic acid. The elution of the gradient began with 5% B at a 200 μL/min flow and increased linearly to 50% B over 7 min, then the elution continued at 50% B for 1.5 min and finally, the column was re-equilibrated for 2.5 min. MS data were acquired in full-scan mode at high resolution on a hybrid Orbitrap Elite (Thermo Fisher Scientific, Bremen, Germany). The system was operated at 240,000 resolution (m/z 400) with an AGC set at 1.0E6 and one microscan set at 10-ms maximum injection time. The heated electrospray source HESI II was operated in positive mode at a temperature of 90 C and a source voltage at 4.0KV. Sheath gas and auxiliary gas were set at 20 and 5 arbitrary units, respectively, while the transfer capillary temperature was set to 275 °C.

Mass spectrometry data were acquired with LTQ Tuneplus2.7SP2 and treated with Xcalibur 4.0QF2 (Thermo Fisher Scientific). Lipid identification was carried out with Lipid Data Analyzer II (LDA v. 2.6.3, IGB-TUG Graz University) (Hartler et al., 2011). The LDA algorithm identifies peaks by their respective retention time, m/z and intensity. Care was taken to calibrate the instrument regularly to ensure a mass accuracy consistently lower than 3 ppm thereby leaving only few theoretical possibilities for elemental assignment. Lipid levels were calculated by normalization over internal standards. Enrichments in viral particles over cells were calculated according to the formula: (viral_Lipid x / viral_PtdCho 34:1)/ (cellular_Lipid x / cellular_PtdCho 34:1)

#### Cholesterol analysis

To detect free cholesterol and cholesteryl esters in samples we used a fluorometric method based on an enzyme-coupled reaction, Cholesteryl esters are hydrolyzed by cholesterol esterase into cholesterol, which is then oxidized by cholesterol oxidase to yield H2O2: Amplex Red Cholesterol Assay kit (Thermofisher).

#### Automated Microscopy analysis of Filipin staining

Vero cells were plated at 10000 cells per well and transfected, with plasmids expressing HA-tagged Spike (WT or 10C-A), for 24 h using TransIT-X2® Transfection Reagent (Mirrus) in Ibidi 96-well µplates. Cells were fixed with 3% PFA and permeabilized with 0.05% saponin. Anti-HA antibody (Roche Diagnostics) was used at 1:100 together with RNase (Qiagen) for primary staining. After washes with automated plate washer (BioTek EL406), cells were incubated for half an hour with Alexa488 anti-Rat (1:200 Jackson Immunoresearch) and filipin diluted 1:50 (Sigma), nuclei were stained by PI (1:200; Thermo-Fisher Scientific). Imaging was acquired on a Molecular Devices™ IXM-C with a 40x plan Apo objective, 49 images were acquired in each well. We had 8 replicas for each condition and for each of 3 independent experiments.

To analyze and quantify free cholesterol (filipin) staining, we used the MetaXpress Custom Module editor software from Molecular Devices, as in previous publications (Larios et al., 2020; Moreau et al., 2019). We first segmented each individual nucleus using the PI channel, we then used the HA channel to mask the expressing cells. The final masks were applied to all original fluorescent images and measurements per cell (nucleus) and average per well were extracted. The same analysis pipeline was applied to all images.

#### Molecular Modeling and Simulations

The model of trimeric Spike’s TM helix + C-terminal region (residues 1201-end in Uniprot P0DTC2) was built by extending laterally the C-terminus of model 1_1_2 built from PDB 6VSB by (Woo et al., 2020). Membrane insertion was carried out with CHARMM-GUI’s web tool in atomistic or coarse-grained (CG) lipid bilayers as required (Hsu et al., 2017; Jo et al., 2008; Qi et al., 2015). For CG molecular dynamics (MD) simulations, we parametrized, also using CHARMM-GUI, the protein with the MARTINI 2.2p CG force field (Marrink et al., 2007), the DPPC, DLiPC and cholesterol with MARTINI CG force field for lipids (Marrink et al., 2004) (notice that DLiPC corresponds to MARTINI’s DIPC lipid), and S-palmitoylated cysteines with the parameters provided by (Atsmon-Raz and Tieleman, 2017). Membranes for CG MD simulations contained 50% DPPC, 30% DLiPC and 20% cholesterol, which we verified forms rafts within 1 μs of CG MD simulation without proteins inserted, as shown by (Lorent et al., 2017). For atomistic MD simulations, we parameterized the protein, POPC membrane and palmitoylated cysteine with CHARMM36m as implemented in CHARMM-GUI. All MD simulations were run in Gromacs (Abraham et al., 2015) using standard settings from CHARMM-GUI for CG and atomistic membrane simulations, at 303 K. In atomistic MD we used a 12 Å cutoff for nonbonded interactions, PME, NVT equilibration followed by NPT production with semi-isotropic pressure coupling to 1 atm and 2 fs timestep; in CG MD we used standard MARTINI equilibration for membranes followed by NPT with semi-isotropic pressure coupling to 1 atm and 20 fs timestep. Simulations were visualized in VMD (Humphrey et al., 1996) and quantitative analysis were carried out through custom VMD Tcl and bash scripts. Membrane perturbation maps were built with MEMBPLUGIN for VMD (Guixà-González et al., 2014).

## SUPPLEMENTARY INFORMATION

**FIGURE S1.**
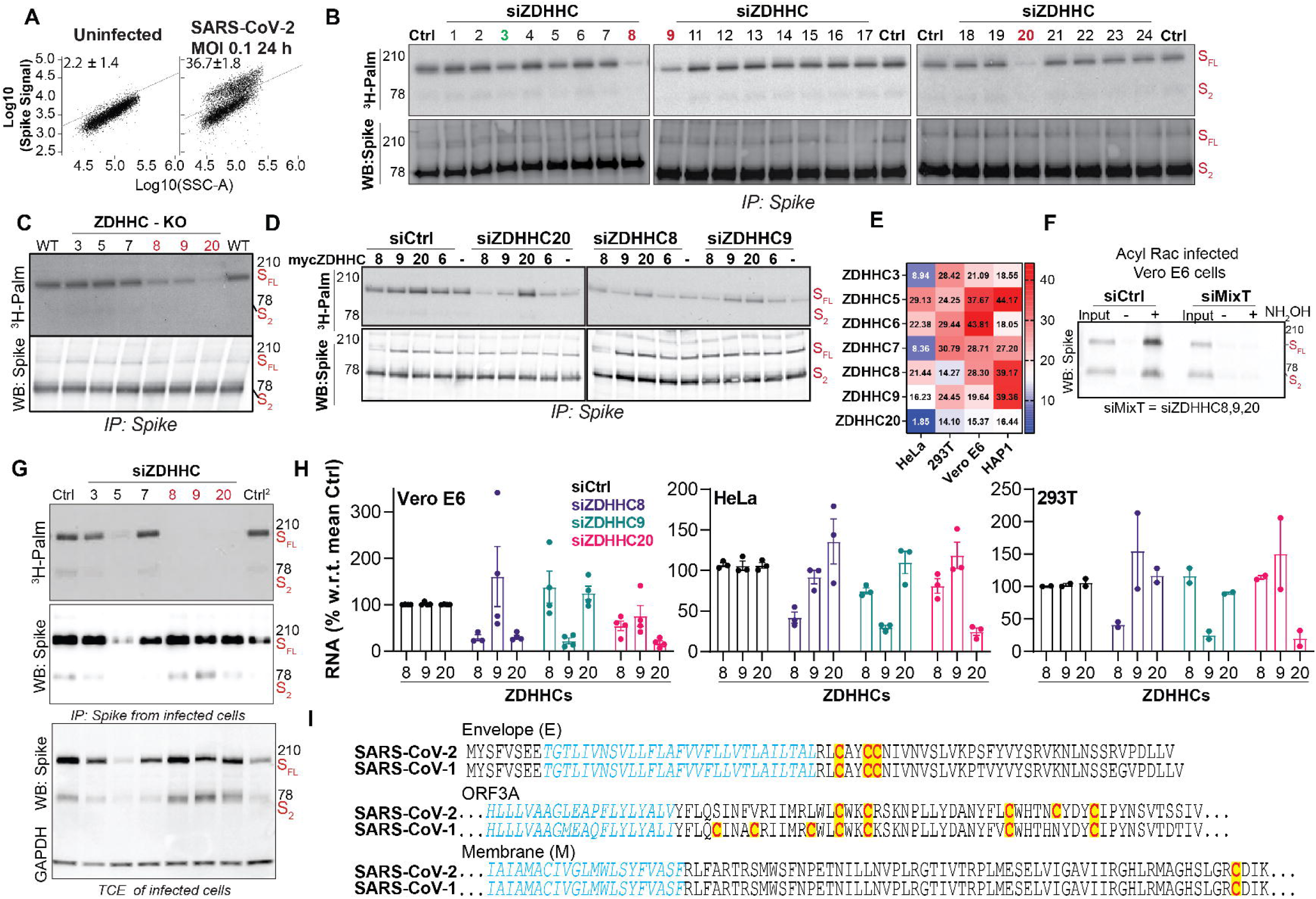
Complement to Figure 1. **A.** Flow cytometry analysis of Vero E6 infected with SARS-CoV-2 (MOI=0.1; 24 h) fixed and stained with Spike antibodies. Dual-parameter density plots (Spike signal vs side scatter SSCA) depicting high-spike positive cells quantified for 3 independent biological replicates. Uninfected cells were used as control. Results are mean±SEM **B-D.** Western blot and autoradiographic analysis of Spike IP fractions from: **B,** Hela cells siRNA-depleted for the indicated ZDHHCs or transfected with control siRNAs (Ctrl), and expressing Spike as described and quantified in Figure 1E; **C,** from HAP-1 KO cells expressing Spike described and quantified in Figure 1F; or **D** from HeLa cells processed as in **B** but co-transfected with the siRNA resistant plasmids expressing myc-tagged ZDHHC8, 9, 20 or 6. In (-) cells were transfected with empty plasmid. Screening experiments were carried at 48 to 72 h of Spike expression, with Spike cleavage being more pronounced at later times. Yet, following a 2 h 3H-palmitate pulse, the majority of labeled Spike was full-length. **E.** ZDHHC mRNA levels of in different cell lines. Values are TAG per Million (TPM) corresponding to 10*6 X ((reads mapped to transcript/transcript length)/ SUM (reads mapped to transcript/transcript length)). Results are mean, n ≥ 3 independent experiments. **F.** Acyl-RAC capture assay in Vero E6 cells transfected (72 h) with an siRNA pool to silence ZDHHC8, ZDHHC9 and ZDHHC20 (siMixT) or siRNA control (Ctrl) and subsequently infected with SARS-CoV-2, MOI≈0.1, 24 h. Cell lysate fractions (input), S-palmitoylated proteins (detected after hydroxylamine treatment, +NH_2_OH) and control fractions (-NH_2_OH) were analyzed by western blot against Spike. **G.** Western blot, against Spike and GAPDH, and autoradiographic analysis of total cell extracts (TCE) or Spike IP-fractions from Vero E6 cells transfected with siRNAs targeting individual ZDHHCs or control siRNA (siCtrl) and infected 24 h with SARS-CoV-2, MOI=0.1. Cells were metabolically labeled throughout the 24 h of infection with ^3^H-palmitic acid and analyzed for Spike S-palmitoylation quantified in Figure 1G. **H.** Q-PCR analysis of mRNA levels in Vero E6, HeLa and HEK293T cells transfected 72 h with Control (siCtrl) or the indicated siRNAs. Results are mean ± SEM, and each dot represents and independent experiment n ≥3. **I.** Alignment of the juxtamembranenous regions of the SARS-CoV-1 or CoV-2 proteins. Transmembrane domains are shown in blue and cytosolic cysteine residues highlighted in yellow. Sequences were retrieved from UNIPROT: Orf3a CoV-2 (P0DTC3), CoV-1 (J9TEM7); E CoV-2 (P0DTC4), CoV-1 (P59637); M CoV-2 (P0DTC5), CoV-1 (P59596).

**FIGURE S2.**
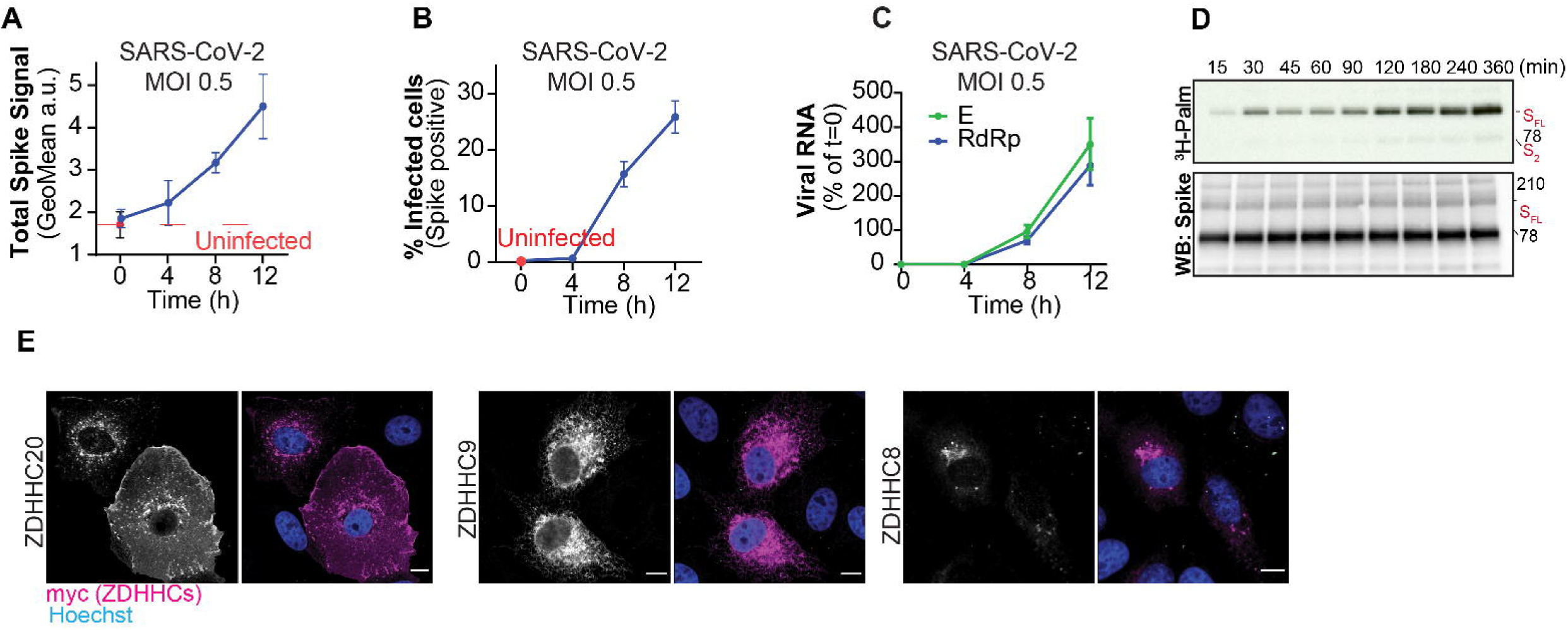
Complement to Figure 2. **A**.**B.C.** Vero E6 were left uninfected (red line) or infected with SARS-CoV-2 MOI≈0.5/1 for the indicated time points. Cells were either harvested and processed for flow cytometry with Spike antibodies (**A-B**) or processed for quantification of total E and RdRp Viral RNA by Q-PCR. Histograms show **(A)** average Spike signal per cell (GeoMean a.u.) **(B)**; % of high-Spike positive cells and **(C)** total viral RNA (E and RdRp) levels in cells throughout infection, quantified for 3 independent biological replicates. Results are mean ±SD. **D.** Representative autoradiography (^3^H-Palm) images and western blot analysis of ^3^H-Palmitic acid incorporation by Spike WT transfected in HeLa cells as described in Figure 2F.**E.** Confocal images of Vero E6 cells ectopically expressing the indicated myc-ZDHHC enzymes (24 h) fixed, and immunolabelled with myc antibodies. Hoechst was used for nuclear counterstaining. scale bar 10 µm.

**FIGURE S3.**
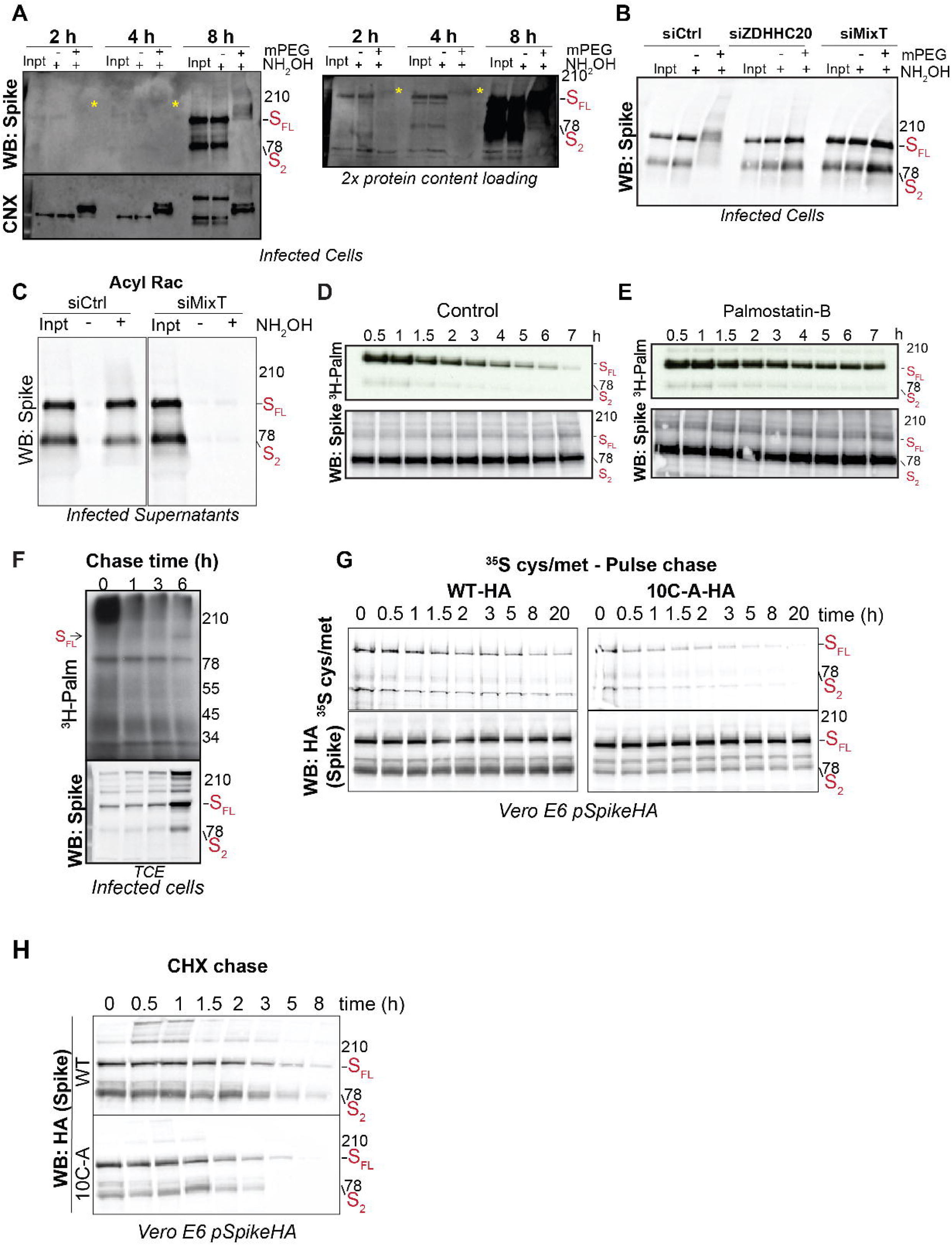
Complement to Figure 3. **AB.** Acy-PEG exchange assay in lysates from Vero E6 cells infected with SARS-CoV-2, **(A)** at high MOI (0.5 to 1) and harvested at indicated time-points or **(B)** transfected for 72 h with the indicated siRNAs (siCtrl; siZDHHC20, or combine ZDHHC8, ZDHHC9, ZDHHC20-siMixT) and infected for 24 h (MOI=0.1). S-acylated proteins were labelled with PEG-5KDa (+mPEG) following hydroxylamine treatment (NH_2_OH) and analyzed with control non-labelled (-mPEG), and input cell lysate fractions (input) by western blot against Spike or **(A)** Calnexin as positive control. In **(A)** a parallel analysis (right panel) was performed using double of the protein content. Yellow asterisks indicate shifted Spike bands at 2 and 4 h p.i. **C.** Acylrac capture assay from pre-cleared, filtered supernatants of Vero E6 siRNA-depleted and infected as in **B**. Hela cells transfected for 24 to 48 h with plasmids expressing untagged Spike were left untreated or pretreated with Palmostatin B for 4 h in serum free medium. Cells were metabolically labeled for a 2 h pulse with ^3^H-palmitic acid, and chased in complete medium for the indicated times. Palmostatin B was maintained throughout the experiment. Levels of incorporated radioactivity in Spike-IP fractions were analyzed by autoradiography (^3^H-Palm) and western blot against Spike. Similar analysis was used to quantify the data depicted in Figure 3E. **F.** Autoradiography (^3^H-Palm) images and western blot analysis of total cell lysate fractions described and quantified in Figure 3G **G.** Representative Autoradiography images and **GH** western blot analysis of: **(G)** ^35^S-Met/Cys pulse chase; and **(H)** Cyclohexamide (CHX) pulse experiments described and quantified in Figure 3H and 3K.

**FIGURE S4.**
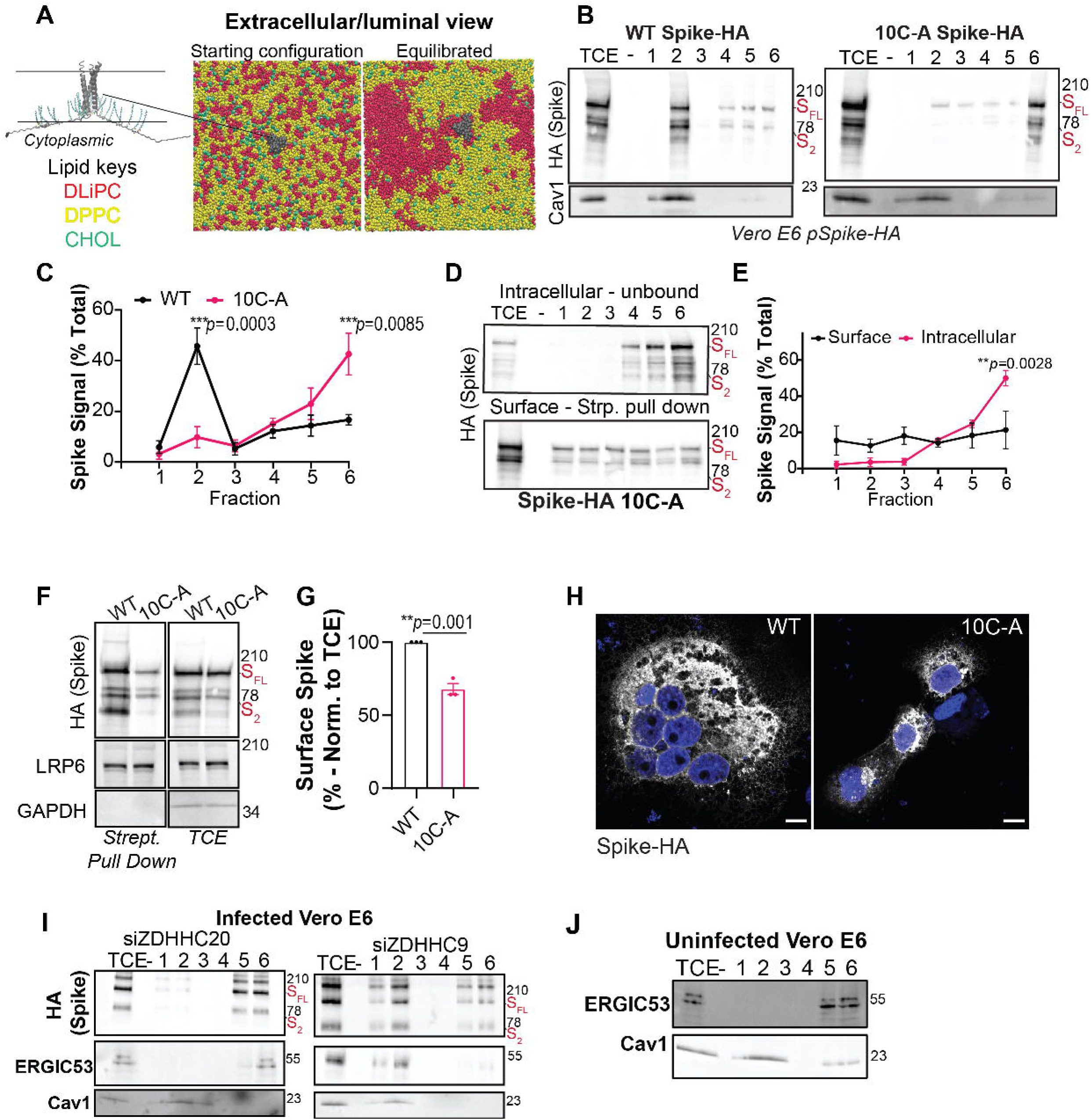
Complement of Figure 4. **A.** (Left) Starting model used for MD simulations shown in atomic detail and with cysteines S-palmitoylated. (Center and Right) correspond to Left and Center of Figure 4A in the main text, but seen from the external side. **B.** Fractionation and DRM isolation (see experimental procedures) of Vero E6 cells expressing HA-tagged Spike (WT or 10C-A mutant – 24 h). Fractions were analyzed by western blot with anti HA (Spike) and Caveolin1 antibodies. **C.** Quantification of Spike-HA (all forms) in each fraction divided by the total signal in all the 6 fractions. Results are mean ± SD, n = 3 independent experiments. **D.** Surface biotinylation and DRM fractionation in Vero E6 cells expressing HA-tagged Spike 10C-A mutant processed as in Figure 4CD. **E** Quantification of HA-Spike signal in each fraction as in D. **FG** - Surface biotinylation assay as in **C** and **G** quantification of total surface proteins pulled-down by streptavidin-beads and analyzed, with correspondent total cell lysate fractions (TCE), by western blot against HA (HA Spike), LRP6 (surface control) and GAPDH (negative control). **H.** Values for Spike surface expression is shown divided by the amount of Spike in Total Cell Extract (TCE). Results are mean± SEM and each dot represent an independent experiment, n=3. **I.** Confocal images of Vero E6 cells transfected as in **B** immunolabelled for HA-Spike, and stained with Hoechst (nuclei). Scale bar-10 µm. **IJ** Western blot analysis of fractions from: **I** Vero E6 cells transfected with siZDHHC20 or siZDHHC9 (72 h) and infected with SARS-CoV-2, 24 h MOI ≈0.1 or **J** uninfected cells Vero E6 cells, processed as in **C**. Western blots against Spike and **JK** ERGIC53 and Cav-1 were used for quantification in Figure 4LM.

**FIGURE S5.**
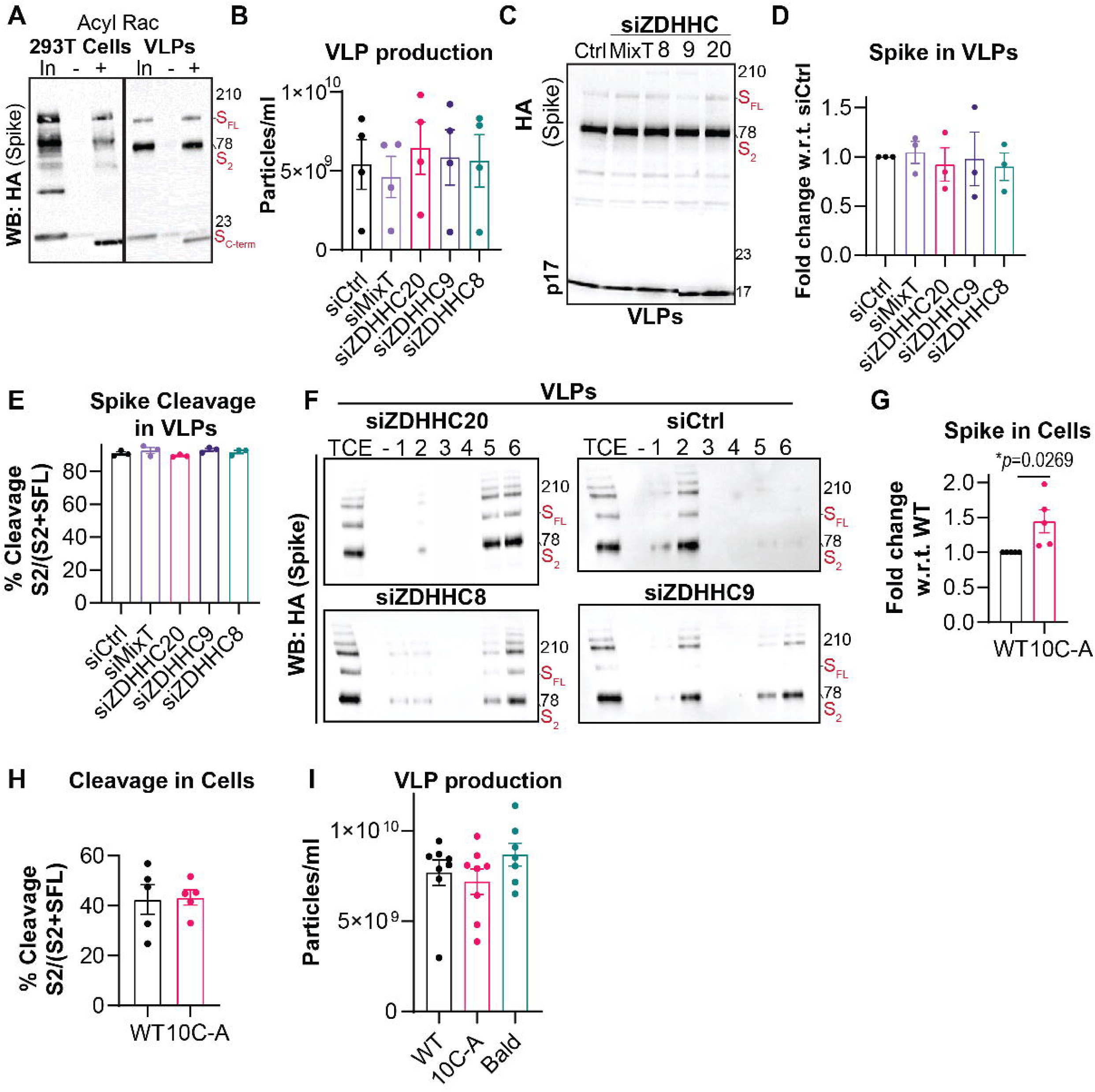
Complement of Figure 5. **A.** Acyl-RAC capture resin assay using HA-Spike-pseudotyped VLPs and correspondent HEK293T producer cells. Cell lysate fractions (input), S-palmitoylated proteins (detected after hydroxylamine treatment, +NH_2_OH) and control fractions (-NH_2_OH) were analyzed by western blot against Spike. **B.** Concentration of VLP suspensions generated in Figure 5A**-C**, measured by ELISA for the lentiviral protein p24. Results are mean ± SEM, and each dot represents an independent preparation. **C, D, E**. Western blot analysis, and quantification of VLPs generated in Figure 5A**-C**. Blots were analyzed for HA Spike, and p17 lentiviral matrix protein. **D.** HA-Spike levels were normalized against p17 levels and set to 1 for WT. **E** Spike cleavage was quantified as the ratio between S2 and S2+S full length. Results are mean ± SEM, and each dot represents an independent VLP preparation **F.** Western blot analysis of fractionation profiles from VLPs generated as in **B** and used for quantification in Figure 5C. **G, H.** Quantification of HA-Spike levels and ratio between S2 and S2+S full length bands (Cleavage) in HEK293Tcells corresponding to the representative blots showed in Figure 5D. *p* values were obtained by students t test **I.** Concentration of non-typed, bald, or HA-Spike (WT or 10C-A mutant) pseudotyped VLP suspensions generated as in Figure 5D**-J**, measured by ELISA for the lentiviral protein p24. Results are mean ± SEM, and each dot represents an independent preparation.

**FIGURE S6.**
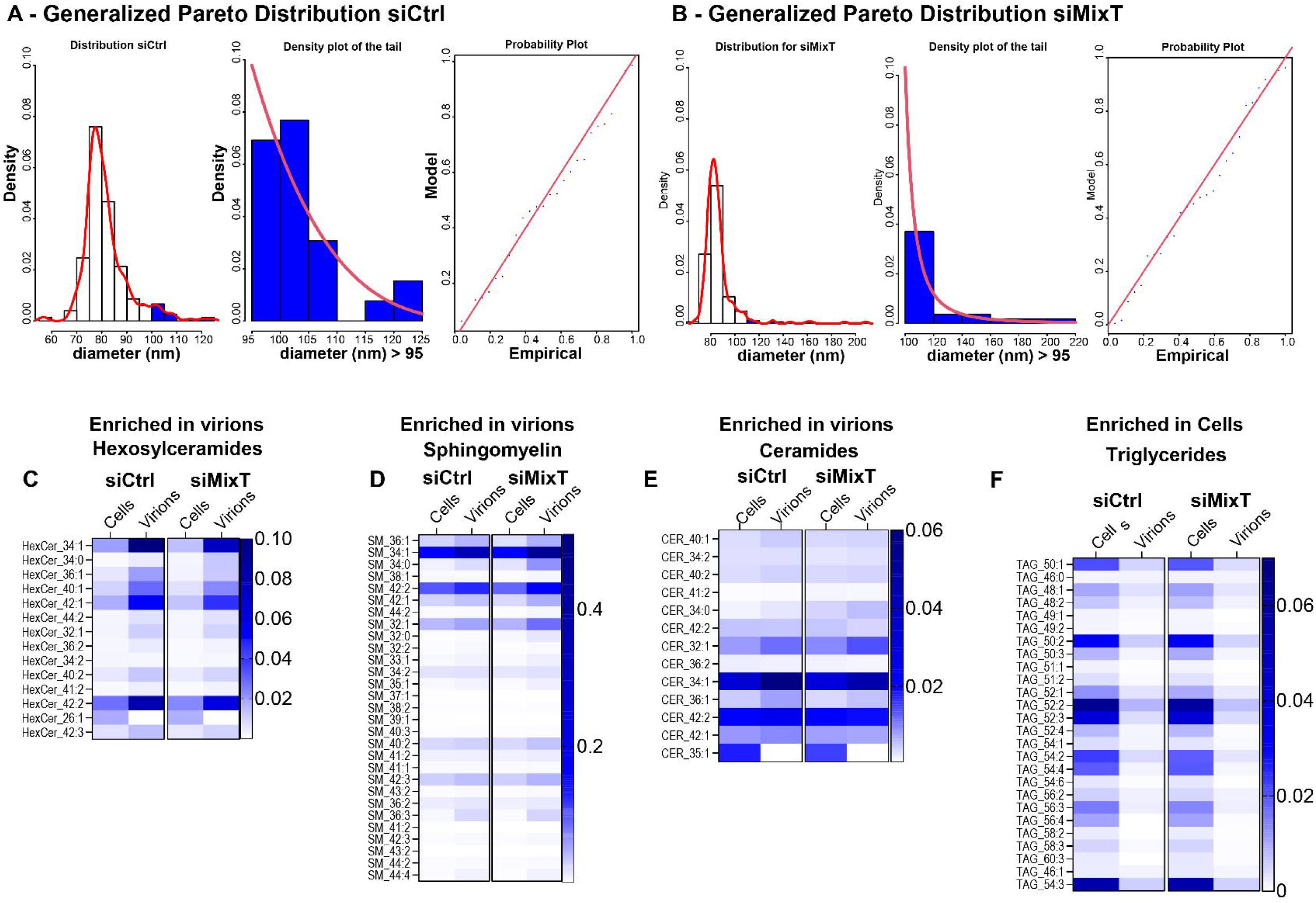
Complement Figure 6. **A-B.** General Pareto (GP) distribution analysis for virion diameters derived from siCtrl or siMixT depleted cells. **left** histograms show Frequency distribution of virion diameters (Bin width 5 nm), with tails distribution in blue used to model the GP distribution. **Centre** histograms show the model for the extreme values of each data set. **Right** graphs show the fitting of the GP distribution analysis to each data set. The theoretical distribution of the extreme values consists in the GP distribution, which can be defined by two parameters: Shape (**ξ**) and scale (β). The shape parameter indicates the propensity of a population to generate extreme values, where negative **ξ**values indicate that a population does not generate extreme values, whereas positive values indicate the propensity of a population to generate extreme limitless values. For siCtrl, **ξ** = −0.20 ±0.19, significantly lower than for siMixT **ξ** = 0.64 ±0.34. **C**-**F**. Heat maps showing an equivalent variation of the levels of individual lipid species from different Lipid classes in SARS-CoV-2 virions derived from siCtrl-treated cells or siMixT-(siZDHHC8/9/20 siRNA pool)-treated cells**. C, D, E** show membrane lipids enriched in SARS-CoV-2 virions **C**, Hexosylceramide (HexCer), **D**, Sphingomyelins (SM) **E,** Ceramides (CER), whereas **F,** show heat maps for non-membranous Triglycerides (TAG) species not abundant in purified virions. Lipid nomenclature is presented as: Lipid class abbreviation (e.g. HexCer), and variable components (fatty acids, Number of C atoms:number of double bonds – e.g. HexCer_34:1).

**FIGURE S7.**
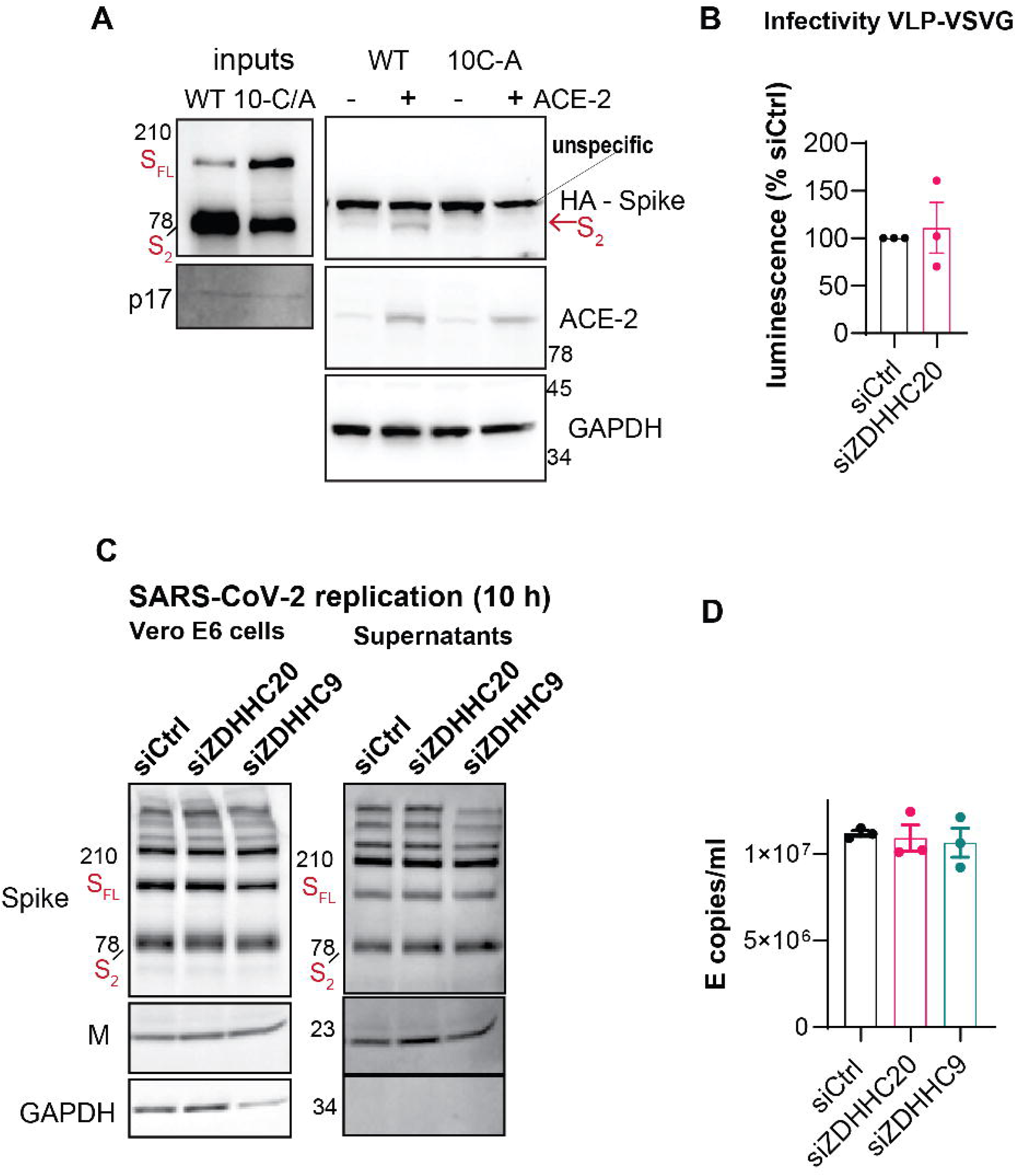
Complement to Figure 7. **A.** Western blot analysis of inputs and total cell extracts from synchronized fusion assays (see experimental procedures) using HeLa or ACE-2-HeLa cells. Surface attached particles were removed by acid wash and internalized Spike S2 signal quantified by western blot against Spike, and expressed as fraction of WT-pseudotyped VLP signal in HeLapACE-2 cells **(see** Figure 7B). The increased exposure of western blot membranes to retrieve signal from the internalize S2 fragment originated the appearance of an unspecific band indicated in the gel. **B** Control infectivity assays as done as in Figure 7D using VSVG-pseudotyped VLPs suspensions harvested from siCtrl or siZDHHC20-silenced cells adjusted to the lentiviral p24 content. **C** Western Blot analysis, probing for viral proteins Spike and M and host cellular GAPDH, from infected total cell extracts (Vero E6 cells) and correspondent supernatants retrieve after 10 h of infection with SARS-CoV-2 at high MOI (1-2) as described in Figure 7G and used to re-infect cells as described in Figure 7H**. D.** Supernatants described in **C** were adjusted to equivalent levels of Viral E RNA copies per ml quantified for three independent preparations.

**Supplentary table 1:**
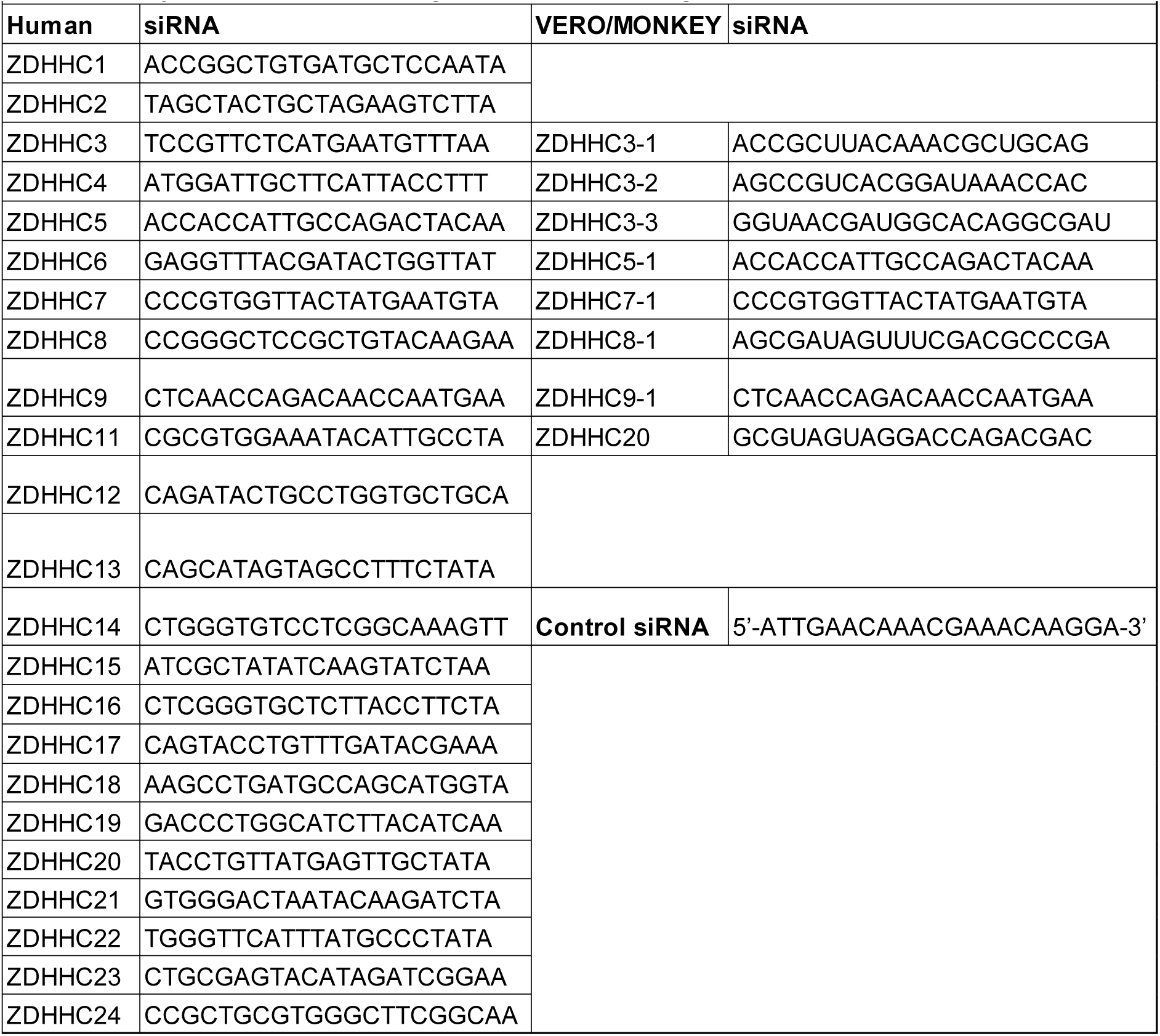
siRNA oligos, related to Oligonucleotides and Plasmids.

**Supplentary table 2:**
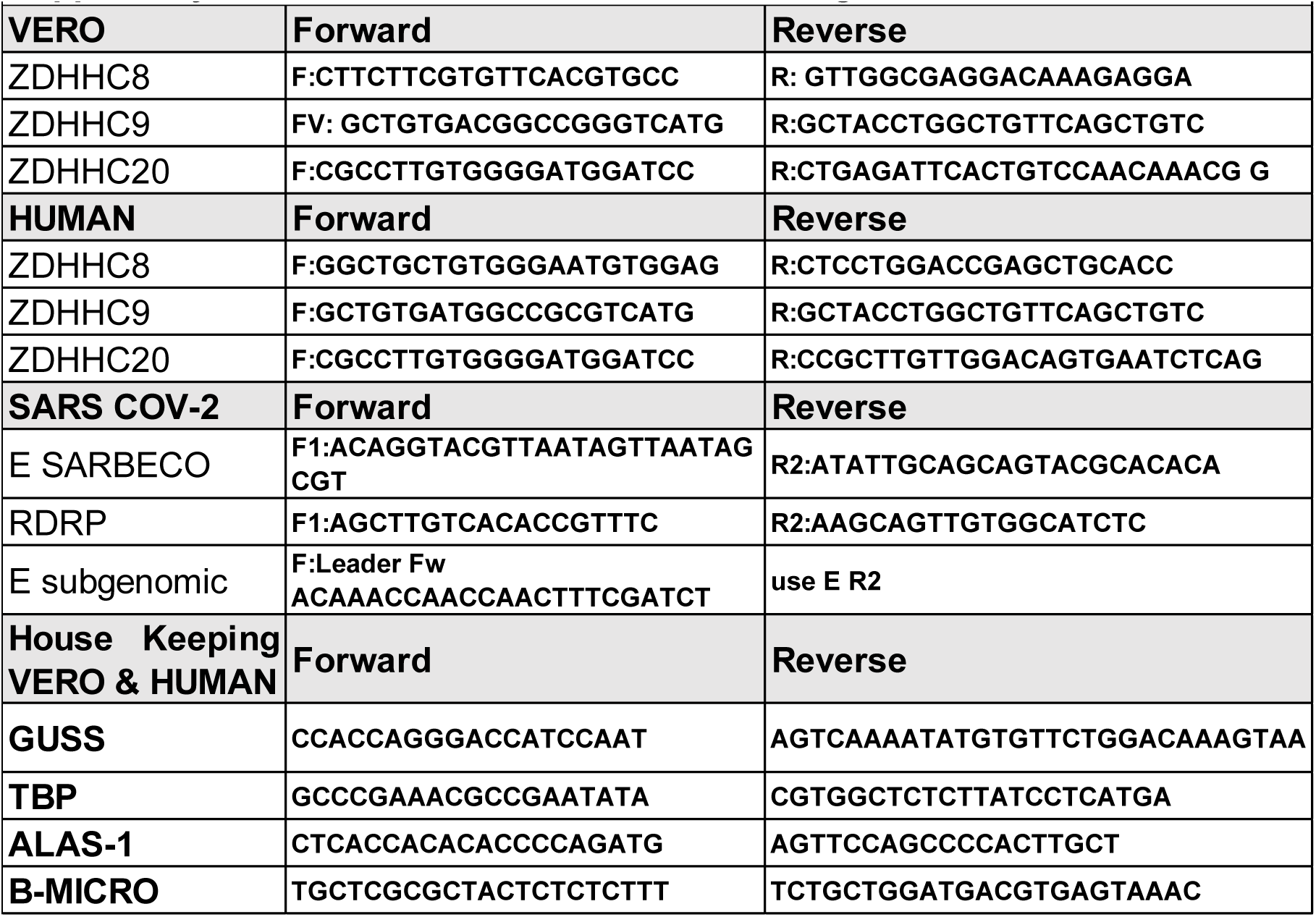
QPCR-Primers, related to Oligonucleotides and Plasmids.

